# Understanding bottlenecks in the microbial production of partially acetylated chitooligosaccharides

**DOI:** 10.1101/2025.02.24.639843

**Authors:** Sofie Snoeck, Hamza Malik, Wouter Demeester, Dries Duchi, Wies Lips, Chiara Guidi, Marjan De Mey

## Abstract

Chitooligosaccharides (COS) are versatile biomolecules with applications across food, pharmaceutical, and cosmetic industries. Expanding the COS portfolio, particularly with partially acetylated COS (paCOS) of defined degree and pattern of acetylation, is essential to unlocking their full potential. This study investigates the co-expression of chitin deacetylases (CDAs) with a chitooligosaccharide synthase (CHS) *Rh*NodC in *E. coli* for *in vivo* paCOS production. While this approach shows promise, it is hampered by reduced overall (pa)COS yields and incomplete conversion of fully acetylated COS to paCOS. Our findings reveal that *Rh*NodC and CDAs co-localize, suggesting potential interactions that influence production efficiency. Additionally, CDA expression induces significant stress responses, including upregulation of *ibp*A and *cpx*P promoters linked to inclusion body formation and membrane stress, respectively. This is accompanied by pronounced cellular elongation, further indicating cellular distress. These bottlenecks highlight the need for deeper exploration of *Rh*NodC-CDA interactions and stress mitigation strategies to optimize scalable *in vivo* paCOS production.

**Highlights:**

‒ Co-expression of rhizobial NodC and chitin deacetylases reduces chitooligosaccharide yield.
‒ Conversion of COS into paCOS remains incomplete upon co-expression.
‒ Upregulation of stress responses suggests protein misfolding and membrane stress.
‒ NodC and chitin deacetylases possibly co-localize and affect cellular localization.
‒ Chitin deacetylase expression causes cell elongation up to 30 micrometers.
‒ Further study needed on rhizobial NodC-chitin deacetylase and substrate interactions.

## 1. Introduction

Chitooligosaccharides (COS) are a class of molecules with a diverse set of biological activities, making them valuable in industries such as food [1,2], feed [3–5], pharma [6–8] and cosmetics [9]. These molecules, composed of β-(1,4)-linked *N*-acetylglucosamine (GlcNAc, A) and/or glucosamine (GlcN, D) residues, are abundant in nature as they are present in fungal cell walls, crustacean shells and insect cuticles [10]. COS owe their biological activity to three structural parameters: degree of polymerization (DP), degree of deacetylation (DDA) and pattern of acetylation (PA) [11]. These parameters influence interactions with biological receptors, following a lock-and-key mechanism that governs COS functionality [12]. This lock-and-key mechanism has been demonstrated in several studies, such as the influence of DA and PA on the priming activity of chitosan tetramers in rice cells, where specific PA variations significantly affect activity [11]. Furthermore, COS has been reported to have a high antioxidant activity, which has been found to vary depending on the DP [13]. Additionally, plant defense responses have been found to peak with COS of DP 6 and DP 7 in *Arabidopsis thaliana* seedlings and *Citrus erythrosa* leaves, respectively [14,15]. These findings demonstrate that the availability of COS products with well-defined DP, DA and PA is essential to unlock their full application potential.

Current COS production methods primarily involve the chemical, mechanical or enzymatic degradation of chitin and chitosan or the chemical synthesis of COS [16,17] (**Figure 1**, left). However, the main drawback of degradation methods is that they yield heterogenous mixtures of (pa)COS molecules. Additionally, the current methods consistently suffer from inefficiencies, batch-to-batch variations, scalability issues, and environmental concerns [18]. *In vitro* enzymatic approaches offer some advantages, such as increased structural control, but remain economically unviable or rely on chemically produced substrates, perpetuating existing challenges [18–22]. Consequently, the reliance on poorly defined COS mixtures limits the study of structure-function relationships, curbing the development of advanced COS applications [16].

**Figure 1:**
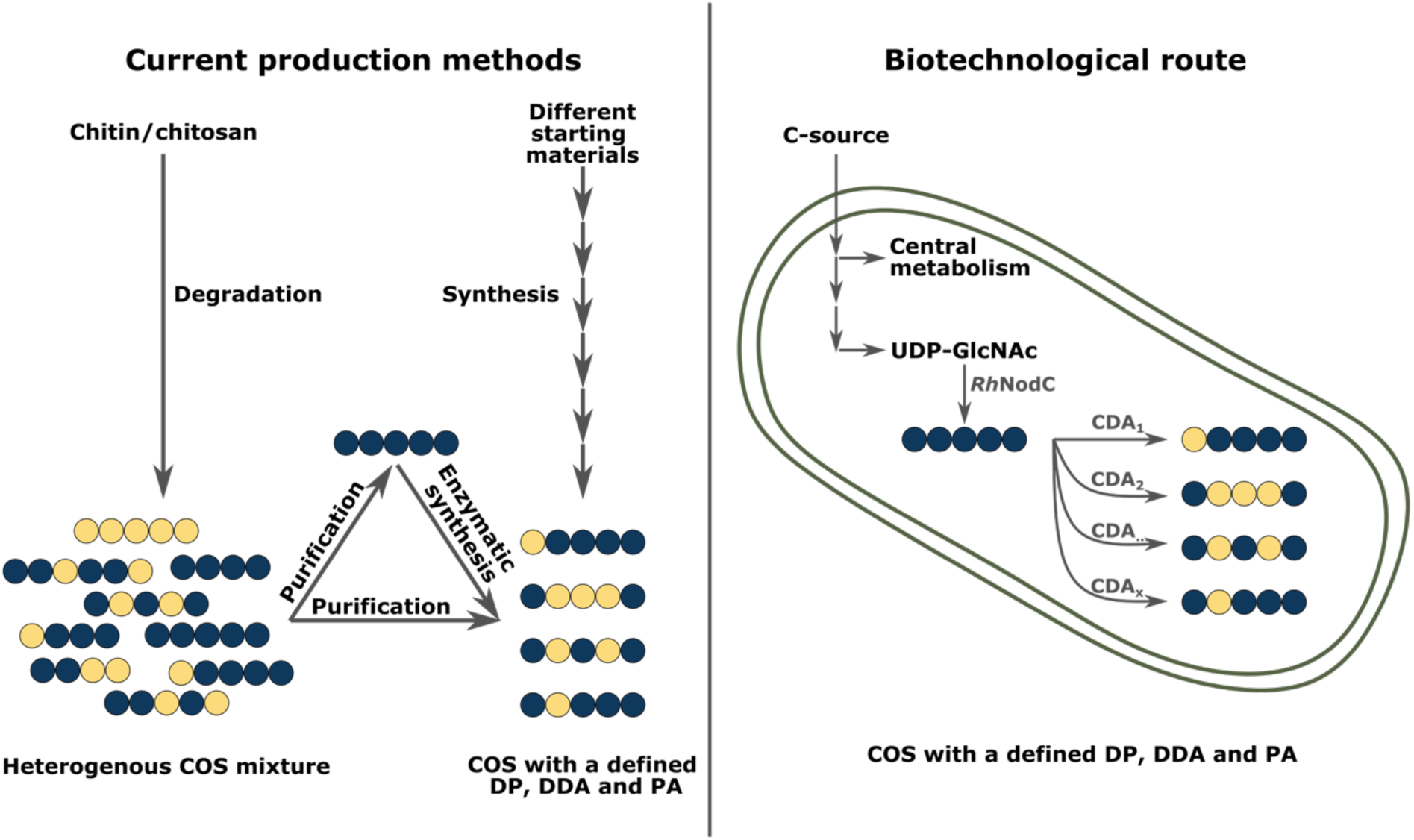
Overview of paCOS production methods. Left: Current production methods relying on the chemical/enzymatic/mechanical degradation of chitin/chitosan into a COS mixture. Defined production is possible via purification or purification and subsequent enzymatic synthesis using regioselective enzymes. Chemical synthesis of defined COS is possible but includes multiple (de)protection steps. Right: Biotechnological production of defined paCOS. The imported carbon source is metabolized towards UDP-GlcNAc, which is converted into A5 by RhNodC. CDAs will take A5 as a substrate and convert it into specific paCOS molecules, depending on their regioselectivity. A5 = fully acetylated chitopentaose; (pa)COS = (partially acetylated) chitooligosaccharides; CDA = chitin deacetylase; GlcNAc = N-acetylglucosamine; DP = degree of polymerization; DDA = degree of deacetylation; PA = Pattern of acetylation; RhNodC = chitooligosaccharide synthase of Rhizobium sp. GRH2; yellow circle = glucosamine; blue circle = N-acetylglucosamine.

A promising alternative is the *in vivo* biosynthesis of COS, leveraging natural enzymatic pathways to produce structurally defined molecules (**Figure 1**, right). The NodC enzyme, a chitooligosaccharide synthase in *Rhizobium* species, synthesizes fully acetylated COS with defined DP, which are then structurally modified by NodB chitin deacetylases (CDAs) through selective deacetylation producing partially acetylated COS (paCOS) [23–26]. Beyond rhizobia, CDAs are widespread in nature—occurring in (marine) bacteria, fungi and insects—where they exhibit diverse regioselectivities that contribute to various biological processes such as fungal pathogenicity, cell wall remodeling, and nitrogen recycling [27,28]. These naturally occurring enzymatic systems provide a foundation for a biotechnological approach to COS production. *In vivo* expression of the *nodC* gene from various rhizobia in *Escherichia coli* (*E. coli*) has already demonstrated successful production of fully acetylated COS [29–31], and co-expression of NodC with CDAs enables the *in vivo* production of paCOS with a defined DP, DDA, and PA [29]. This strategy represents a significant step toward expanding the COS portfolio, particularly in generating tailored paCOS molecules. In recent decades, paCOS have gained increasing interest due to their enhanced solubility and biological activities [32]. Despite the well-characterized *in vitro* properties of many CDAs [33–36], their *in vivo* application remains underexplored [29], posing challenges for optimizing production systems [19,28].

The underexplored nature of *in vivo* CDA applications is largely due to the considerable difficulties associated with *in vivo* expression, such as growth retardation and functionality loss. These challenges underscore the need for a more thorough investigation into the constraints of *in vivo* CDA expression, as a deeper understanding of these factors will be critical for optimizing production systems and enabling successful paCOS synthesis on a larger scale (**Figure 1**, right). Hence, this research investigates how CDA expression impacts cellular stress responses in *E. coli* by monitoring the activation of specific stress pathways, while exploring the subcellular localization of *Rh*NodC and CDAs using microscopy to identify potential engineering targets for enhancing paCOS production. By identifying these bottlenecks, this study seeks to enhance the efficiency of *in vivo* paCOS production, paving the way for scalable and economically viable biotechnological processes that unlock the full potential of defined paCOS for industrial applications.

## 2. Results and discussion

### 2.1. *In vivo* production of (partially acetylated) chitooligosaccharides

To demonstrate the challenges encountered during *in vivo* CDA applications, several CDAs were expressed together with the chitooligosaccharide synthase NodC from *Rhizobium sp.* GRH2 (*Rh*NodC), which produces fully acetylated chitinpentaose (A5). CDAs utilize the product of *Rh*NodC as their substrate and deacetylate one or more GlcNAc residues, resulting in paCOS. Specifically, three bacterial CDAs (*Rh*NodB, *Vc*COD, *Bs*PdaC) were selected for *in vivo* testing (Table 1). While fungi also produce CDAs, no fungal CDAs were chosen as working with fungal enzymes *in vitro* and *in vivo* poses significant challenges, primarily due to difficulties in protein expression and purification [37]. In preparation of the experiments, any transmembrane (*Bs*PdaC) or signal peptide regions (*Vc*COD) present in the CDAs were removed to avoid any membrane stress related responses.

**Table 1:** Chitin deacetylases used in this study, their organism of origin and their characterized pattern of acetylation (in vitro). CDA = chitin deacetylase; A/GlcNAc = N-acetylglucosamine; D = glucosamine; Ax/Dx = x indicates the number of residues; DP = degree of polymerization; Rh = Rhizobium sp. GRH2; Vc = Vibrio cholerae; Bs = Bacillus subtilis.

| CDA | Organism of origin | Pattern of acetylation | References |
| --- | --- | --- | --- |
| *RhNodB* | *Rhizobium* sp. GRH2 | DA, DA2, DA3, DA4, DA5 | [18,24,25] |
| *VcCOD* | *Vibrio cholerae* | AD, ADA, ADA2, ADA3, ADA4 | [18,38] |
| *BsPdaC* | *Bacillus subtilis* | For DP = 3-5: Starting from the middle residue(s) sequentially deacetylates all GlcNAc residues, except for the reducing end | [39,40] |

Initially, a production assay was performed to benchmark the functionality of the CDAs *in vivo*. Both *Rhnod*C and the CDAs were expressed from a single plasmid in a triple knockout (Δ*chb*BCARFGΔ*chi*AΔ*nag*Z) *E. coli* strain (3KO) as background [41]. In the 3KO strain, native *E. coli* genes encoding proteins involved in the degradation and/or modification of COS are knocked out (Table 4). As a negative control, an identical plasmid but with a non-coding sequence (*nonCDS*) was expressed, while a *RhnodC* containing plasmid served as a positive control for A5 production [41]. Strains were grown on minimal medium glucose, growth was followed and (pa)COS samples were taken in the exponential, late exponential and stationary phases (Figure 2). Note that throughout this study, only A5 and monodeacetylated chitopentaose (DP = 5, DDA = 20%) were measured as no standards were available to verify the presence or quantity of other paCOS products. All statistical analyses can be found in Supplementary Tables 2-6.

**Figure 2:**
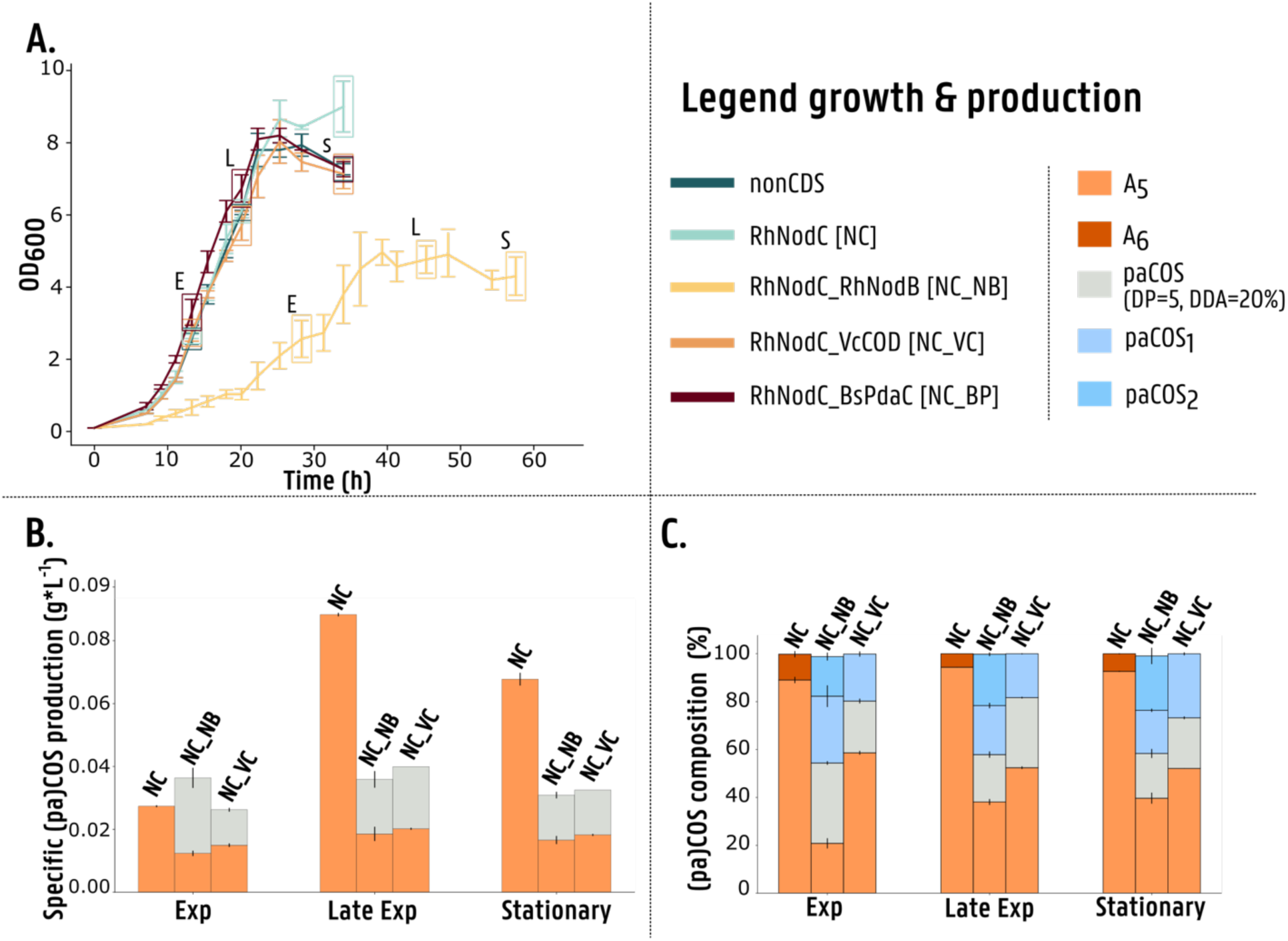
Production assay of paCOS. **A.** Growth curves of the different strains in the production assay. Lines represent the average OD_600_ per time point and error bars represent the standard deviation based on 3 biological replicates. Time points where (pa)COS samples were taken are indicated with a box (E = exponential phase, L = late exponential phase and S = stationary phase). Note that for strain RhNodC-RhNodB sample L is rather in the early stationary phase, while S is in the late stationary phase. **B.** Specific (pa)COS production (g*L^-1^). Bars represent the average production and error bars the standard error of the mean based on three biological replicates. **C.** (pa)COS composition (%). Bars represent ratios of the different (pa)COS that were produced and error bars the standard error of the mean based on three biological replicates. Details on the strains and plasmids used in this study can be found in Table 4 and Supplementary Table 8. Statistical comparisons can be found in Supplementary Tables 2-6. (pa)COS = (partially acetylated) chitooligosaccharides; A5 = fully acetylated chitopentaose; paCOS (DP=5, DDA=20%) = chitopentaose with a degree of deacetylation of 20%; paCOS1 and paCOS2 = unidentifiable paCOS; OD_600_ = optical density measured at 600 nm; RhNodC = chitooligosaccharide synthase from Rhizobium sp. GRH2; RhNodB = chitin deacetylase from Rhizobium sp. GRH2; VcCOD = chitin deacetylase from Vibrio cholerae; BsPdaC = chitin deacetylase from Bacillus subtilis.

Growth of all strains, except 3KO+*Rh*NodC_*Rh*NodB, was similar to the control strains 3KO+*nonCDS* or 3KO+*Rh*NodC (Figure 2 A.). The specific production profile of the strains expressing a CDA differed from the typical patterns observed for COS production (Figure 2 B.). Generally, COS production is growth related and as is the case for 3KO+*Rh*NodC, production increases towards the late exponential phase (0.089 g*L^-1^) and then decreases towards the (late) stationary production phase. Although the 3KO+*Rh*NodC_*Bs*Pdac strain exhibited growth, its specific COS production remained below the detection limit throughout all production phases. Therefore, this strain was excluded from the production graphs. During the exponential phase, total COS production was generally similar to the control strain (3KO+RhNodC) (one-way ANOVA: p = 1.46E-08) while in some cases, COS production was even significantly higher, as observed for 3KO+RhNodC_RhNodB (p = 2.39E-02). However, for all strains co-expressing a CDA, total COS production in the late exponential phase was significantly lower, amounting to about one-third of the production seen in the control strain (3KO+*Rh*NodC). Furthermore, only half of the total COS was converted into paCOS (DP=5, DDA=20%), with the other portion still present as A5, indicating incomplete conversion by the CDAs.

The ratios of (pa)COS (%) over the three production phases confirm that the *Rh*NodC control strain mainly produces A5 (Figure 2 C.). Additionally, in all three phases, a minor fraction of chitinhexaose (A6) production was also noted in the control strain [42]. In strains co-expressing a CDA, additional COS peaks were observed alongside the target paCOS (DP=5, DDA=20%). As COS are reducing oligosaccharides, they appear in either an α-or β-anomeric form and subsequently depict a distinctive double peak due to the limited separation capacity of the HPLC column. In more detail, A5, paCOS (DP=5, DDA=20%) as well as two other unidentifiable COS peaks, termed paCOS1 and paCOS2, were observed. More specifically, the 3KO+*Rh*NodC_*Vc*COD strain only produced A5, paCOS (DP=5, DDA=20%) and paCOS1 while the 3KO+*Rh*NodC_*Rh*NodB strain additionally produced paCOS2. Furthermore, the 3KO+*Rh*NodC_*Rh*NodB strain shows a steady decrease in paCOS ratio throughout all three production phases. This suggests that paCOS molecules are either further converted into non-detectable paCOS products or are subject to degradation over time. Given that in the used 3KO strains, all genes encoding for COS degrading or modifying proteins were removed, the latter should not be the case. A possible hypothesis is that CDAs may also exhibit reverse activity, *N*-acetylating the paCOS back into its fully acetylated form. While this has been demonstrated and validated *in vitro*, it remains unproven *in vivo* [43]. However, this phenomenon could also result from an unidentified mechanism linked to the co-expression of NodC and CDAs.

These results confirm that *in vivo* expression of CDAs is not optimal. The 3KO+*Rh*NodC_*Rh*NodB strain experienced severe stress, leading to impaired growth. For all strains co-expressing a CDA, the conversion of A5 into paCOS (DP=5, DDA=20%) was incomplete. Additionally, the total COS production in the late exponential and stationary phases was significantly lower than that of the control production strain 3KO+*Rh*NodC. The production of both A5 and paCOS (DP=5, DDA=20%) was compromised in the late exponential and stationary phases due to currently unknown reasons. To identify the bottlenecks associated with *in vivo* CDA expression, a comprehensive study was conducted.

### 2.2. Identifying the bottlenecks of the *in vivo* production of (partially acetylated) COS

To unravel the bottlenecks associated with the *in vivo* production of (pa)COS, a stress promoter assay for the activation of various stress mechanisms upon CDA expression was conducted. Furthermore, the cellular localization of the CDAs – with and without *Rh*NodC – was analyzed (Figure 3). The combined interpretation of these assays is important to understand if the presence of *Rh*NodC or one of the COS products influences cell viability.

**Figure 3:**
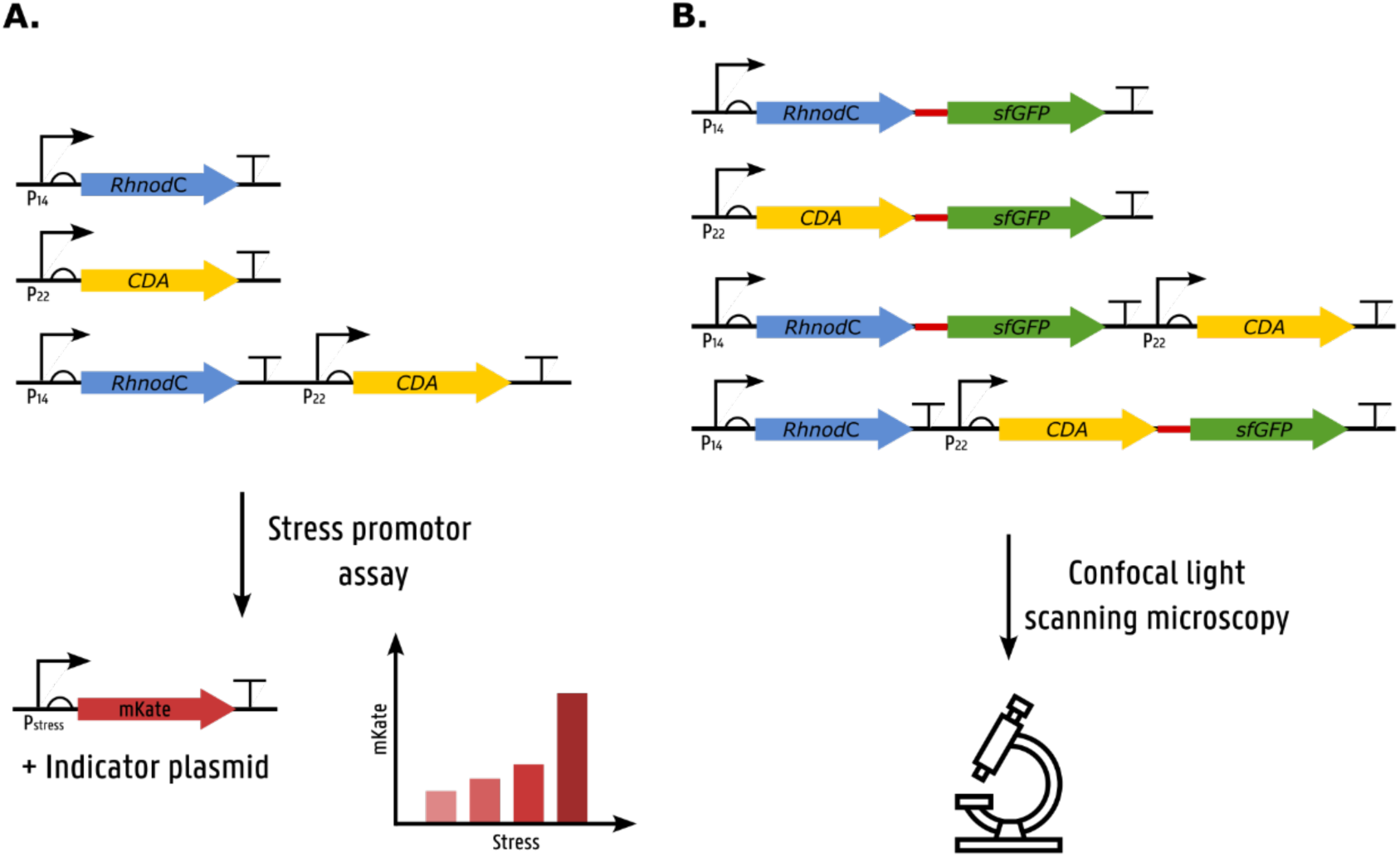
Overview of the studies done to unravel the bottlenecks associated with in vivo CDA expression. **A.** Strains expressing RhnodC and/or the CDAs were supplemented with an indicator plasmid expressing the red fluorescent protein mKate2 under control of different stress promoters (P_stress_) (**Table 2**). Activation of the stress promoters was then assessed by measuring mKate2 intensities. **B.** RhnodC and/or CDAs were linked with the green fluorescent protein sfGFP. These strains were used for confocal light scanning microscopy. More information on the strains and plasmids can be found in Table 4 and Supplementary Table 8. CDA = chitin deacetylase (RhNodB, VcCOD, BsPdaC); Rh = Rhizobium sp. GRH2; Vc = Vibrio cholerae; Bs = Bacillus subtilis; sfGFP = superfolder green fluorescent protein. Gene structures are displayed according to the SBOL guidelines [44].

**Table 2:** Stress promoters tested in the stress promoter assay. The sequences of the promoters can be found in Supplementary Table 1. For more background information on the different stress mechanisms, we refer to [47]. ppGpp = guanosine tetra-and pentaphosphate; ACP = acyl carrier protein; sHSP = small heat shock protein; tRNA = transfer RNA; sRNA = small RNA; σ = sigma factor.

| Promoter | Stress response | Regulation | References |
| --- | --- | --- | --- |
| relA | Stringent response | - *relA*: synthesizes ppGpp<br>- *relA* activated in response to uncharged tRNAs in the ribosomal A-site<br>- Transcriptional activation by different stresses | [50]<br>[51]<br>[52] |
| spoT | Stringent response<br>Nutrient starvation | - *spoT*: synthesizes/hydrolyzes ppGpp<br>- Activity shifts from hydrolysis to synthesis under nutrient starvation (interaction with ACP) | [53]<br>[54]<br>[52] |
| suhB | Stringent response | - *suhB*: Transcription antitermination, promotion of correct ribosome assembly<br>- Transcriptional inhibition by ppGpp | [52]<br>[55] |
| iraP | Stringent response<br>General stress response | - *iraP*: anti-adaptor protein for RssB (adaptor protein for $\sigma^S$ degradation)<br>- Transcriptional activation by ppGpp<br>- Translation dependent on SpoT | [56]<br>[57] |
| dsrA | Stringent response<br>General stress response | - sRNA *dsrA* increases translation of *rpoS* mRNA<br>- Transcriptional activation by DksA and ppGpp | [58]<br>[59] |
| rpoS | General stress response | - Encodes $\sigma^S$, activating > 500 genes<br>- Transcriptional regulation: CRP, arcA, Fis, mqsA<br>- Translationally regulated by sRNA *dsrA*<br>- Post-translationally regulated by ClpXP | [58]<br>[60]<br>[61]<br>[56]<br>[62] |
| ibpA | Heat shock response | - IbpAB: sHSP that co-aggregate with denatured proteins and facilitate refolding<br>- Transcriptional activation by $\sigma^H$<br>- Translational inhibition by IbpA | [63]<br>[64] |
| rpoH | Heat shock response | - Encodes $\sigma^H$, regulates genes for chaperones and proteases, cell homeostasis<br>- Master regulator<br>- Transcriptional regulation: $\sigma^E$<br>- Translational regulation: Temperature, IbpA<br>- Post-translationally regulated by FtsH<br>- Regulation by $\sigma^S$? | [64]<br>[65]<br>[66]<br>[60]<br>[67] |
| cpxP | Membrane stress response | - *cpxP*: CpxP (periplasmic protein) and sRNA *cpxQ*<br>- Transcriptional regulation by cpxR (activated by perturbations in inner membrane & other stresses induced by heterologous protein expression) | [68]<br>[45]<br>[69] |

To this end, two different sets of constructs were made (more detailed information on the promoter and coding sequences can be found in Supplementary Table 1 and the plasmids in Supplementary Table 8). For the stress promoter assay, nine promoters associated with key stress responses commonly triggered during heterologous protein expression were selected. These stress promoters are regulatory DNA sequences that respond to specific cellular stress conditions, such as oxidative stress, protein misfolding, or membrane stress, by activating the transcription of downstream genes. To assess whether a stress mechanism is activated upon *Rh*NodC and/or CDA expression (Figure 3 A.), an indicator plasmid (pInd) was used. This plasmid carries the red fluorescent protein mKate2 under the control of different stress-responsive promoters, allowing fluorescence-based monitoring of stress activation [45]. More information on the selected promoters and their corresponding stress responses is provided in Table 2.

The log_2_ fold changes (log_2_FC) of the corrected fluorescence relative to the control strain (3KO+*nonCDS*+pInd) are given in Supplementary Figure 1B. and Figure 4 (close-up for *ibp*A and *cpx*P). In this experiment, a log₂FC threshold of 1 was set to indicate biological significance, which corresponds to a two-fold increase relative to the control strain. This threshold is commonly used in differential expression studies to identify biologically relevant changes [46]. The general stress (*rpo*S) and heat shock (*rpo*H) responses exhibited only minor effects, with log_2_FC values below 1. However, this result does not rule out the possibility that these responses were not activated, as many of them are regulated on multiple levels (Table 2) [47]. It is possible that transcriptional or translational induction/repression was limited and therefore undetectable in the assay, or that regulation mainly occurs at a post-translational level [48,49].

**Figure 4:**
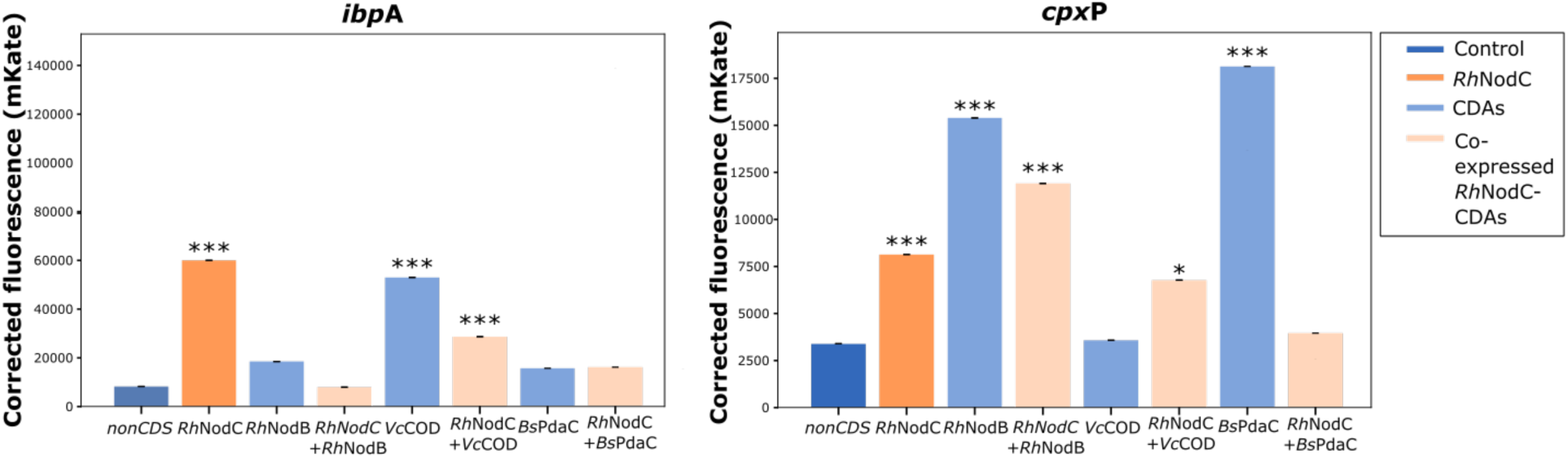
Characterization of the activation of stress promoters in strains expressing RhNodC and/or CDAs (RhNodB, VcCOD and BsPdaC). The corrected fluorescence (mKate2) for the stress promoters ibpA (left) and cpxP (right) is given. Bars represent the mean corrected fluorescence in the stationary phase and error bars represent the standard error of the mean, based on five biological replicates. For statistical comparison of mKate2 levels, one-way ANOVA followed by Tukey HSD for multiple comparison was conducted (all statistical results can be found in Supplementary Table 7). Significance levels compared to the control (3KO+nonCDS+pInd) are represented in the figure. *: p < 0.05; ***: p < 0.001. Details on the strains and plasmids used can be found in Table 4 and Supplementary Table 8. nonCDS = non-coding sequence; CDA = chitin deacetylase; RhNodC = chitooligosaccharide synthase from Rhizobium sp. GRH2; RhNodB = chitin deacetylase from Rhizobium sp. GRH2; VcCOD = chitin deacetylase from Vibrio cholerae; BsPdaC = chitin deacetylase from Bacillus subtilis.

Overall, the stringent response genes, *rel*A, *spo*T, *suh*B, *ira*P and *dsr*A, exhibited a non-significant response compared to the other stress responses. However, a couple of notable exceptions could be observed. *suh*B is negatively regulated by ppGpp and is downregulated in all strains, expect in the 3KO+*Vc*COD strain. In the 3KO+*Vc*COD strain, this promoter was activated with a log_2_FC of 4.6, being the highest log_2_FC recorded in this assay, and growth was halted. However, compared to the normal growth pattern observed in the previous paCOS production assay (Figure 2 A.), these results suggest that the stress promoters—and consequently, the indicator plasmids—could interfere with cellular mechanisms, leading to aberrant outcomes.

Two promoters exhibited upregulation compared to the control: *ibp*A and *cpx*P (Figure 4). More specifically, all strains but 3KO+*Vc*COD and 3KO+*Rh*NodC_*Bs*PdaC showed upregulation for *cpx*P, while all strains but 3KO+*Rh*NodB, 3KO+*Bs*PdaC and 3KO+*Rh*NodC_*Rh*NodB exhibited upregulation for *ibp*A. In general, this upregulation was both biologically and statistically significant. The *Rh*NodC control strain exhibited significant activation of both stress promoters. For the CDAs alone, 3KO+*Vc*COD showed the highest activation for *ibp*A, while for *cpx*P, this was valid for 3KO+*Rh*NodB and 3KO+*Bs*PdaC. Co-expression of *Rh*NodC*_Rh*NodB seemed to reduce activation of the *ibp*A promoter, while for all other combinations the activation remained significant for both stress promoters. The *ibp*A and *cpx*P promoters are connected to inclusion bodies (IBs) and inner membrane stress, respectively. These results suggest that protein misfolding leads to the formation of IBs, or alternatively, the proteins localize within or cause damage to the inner membrane.

To further study the suggested IB and inner membrane stress responses, the strains were analyzed by visualizing the proteins within the cell through confocal light scanning microscopy. For this, superfolder green fluorescent protein (sfGFP) was C-terminally linked to the proteins of interest to minimize interference with potential N-terminal transmembrane or localization signals (Figure 3 B.). Figure 5 shows the results of the confocal light scanning microscopy for *Rh*NodC and *Rh*NodB, all other profiles can be found in the supplementary Figures 2-3. The localization of the proteins and characteristics of all strains are summarized in Table 3. It should be noted that, as previously mentioned, any transmembrane (*Bs*PdaC) or signal peptide regions (*Vc*COD) present in the enzymes were removed to avoid any membrane stress related responses. When expressed separately, *Rh*NodC-sfGFP localized in IBs and *Rh*NodB-sfGFP was found in the cytoplasm. However, when *Rh*NodC was co-expressed with *Rh*NodB, different cellular populations could be distinguished that localize these proteins in either the cytoplasm, membrane or in IBs. Thus, the proteins seem to co-localize. A similar phenomenon was observed for the co-expression of *Rh*NodC with *Vc*COD. In contrast, *Rh*NodC-*Bs*PdaC co-expression does not seem to follow this pattern. For independent expression of *Bs*PdaC, the enzyme was localized in the cytoplasm as well as in the membrane. On the other hand, localization in IBs and the cytoplasm was observed when sfGFP-tagged *Rh*NodC was co-expressed with *Bs*PdaC. When sfGFP-tagged *Bs*PdaC was visualized upon co-expression, the enzyme was only found in the membrane which is unexpected as the transmembrane domain of *Bs*PdaC was previously removed. This does however suggest co-localization, given that *Rh*NodC is a membrane protein. Notably, in nature, *Rh*NodC and *Rh*NodB are transcribed from the same operon, *nod*ABC [70,71]. *Rh*NodC usually localizes to the membrane, and it has been suggested that loose biosynthetic complexes could be formed at the membrane as all other *nod* genes are known to be cytoplasmic [72]. Our results suggest that the chitooligosaccharide synthase *Rh*NodC and some of the CDAs interact with each other. It is unclear whether these interactions are protein-protein interactions (*Rh*NodC with CDA) or whether the CDA interacts with the product of *Rh*NodC, A5. To study possible protein-protein interactions between the enzymes, *Rh*NodC can be rendered non-functional [23,73,74] by mutating the important residues in the catalytic center (D241) [74]. Additionally, interactions could be confirmed using fluorescence energy transfer (FRET) [75].

**Figure 5:**
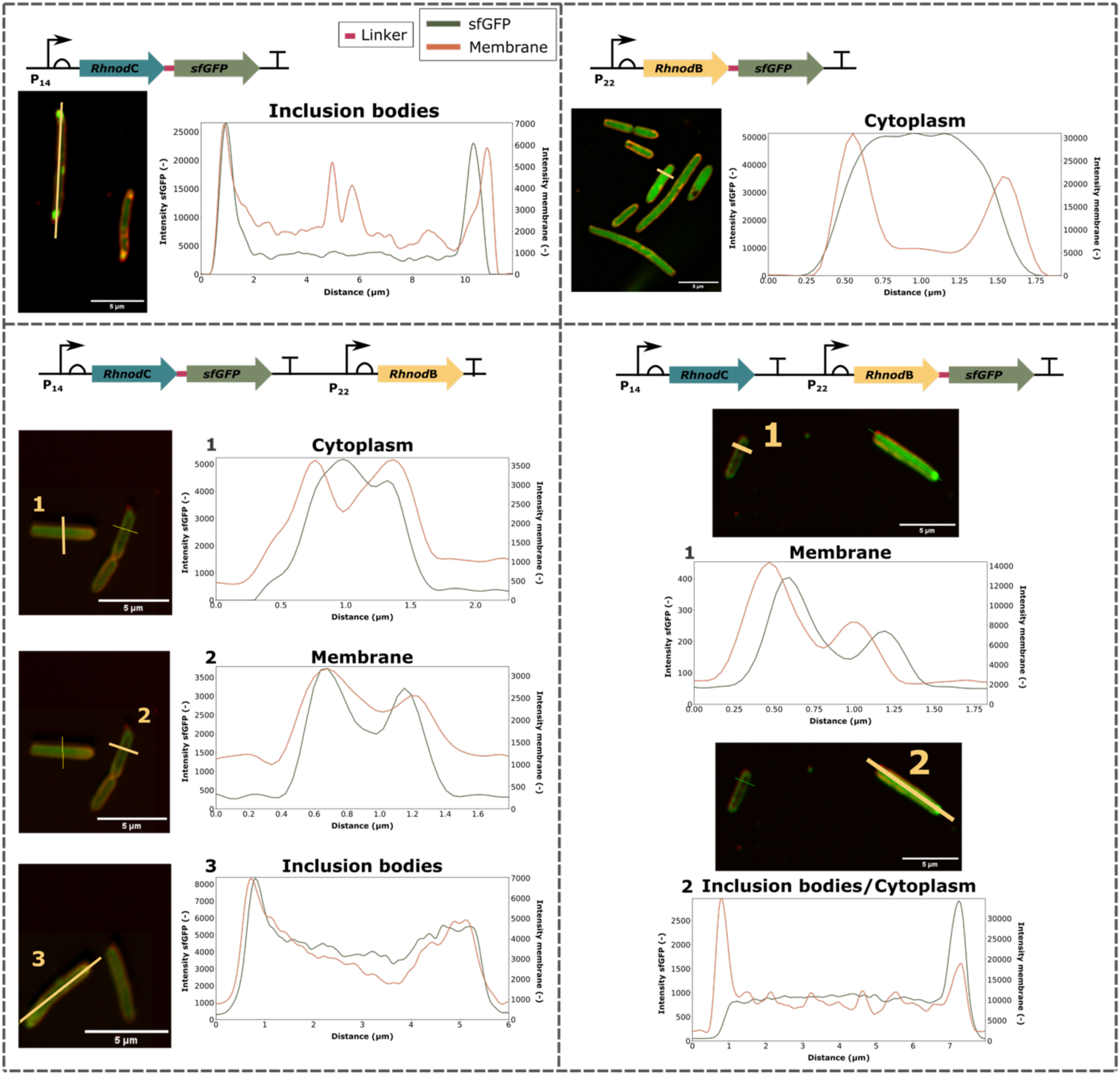
Confocal light scanning microscopy images of RhNodC and the CDA, RhNodB, with intensity cross-section profiles for both the red FM4-64 membrane dye (orange) and sfGFP (green). The cross-sections analyzed are indicated with a yellow line in the images. More information on the strains and plasmids used can be found in Table 4 and Supplementary Table 8. RhNodC = chitooligosaccharide synthase from Rhizobium sp. GRH2; RhNodB = chitin deacetylase from Rhizobium sp. GRH2; sfGFP = superfolder green fluorescent protein. Gene structures are displayed according to the SBOL guidelines [44].

**Table 3:**
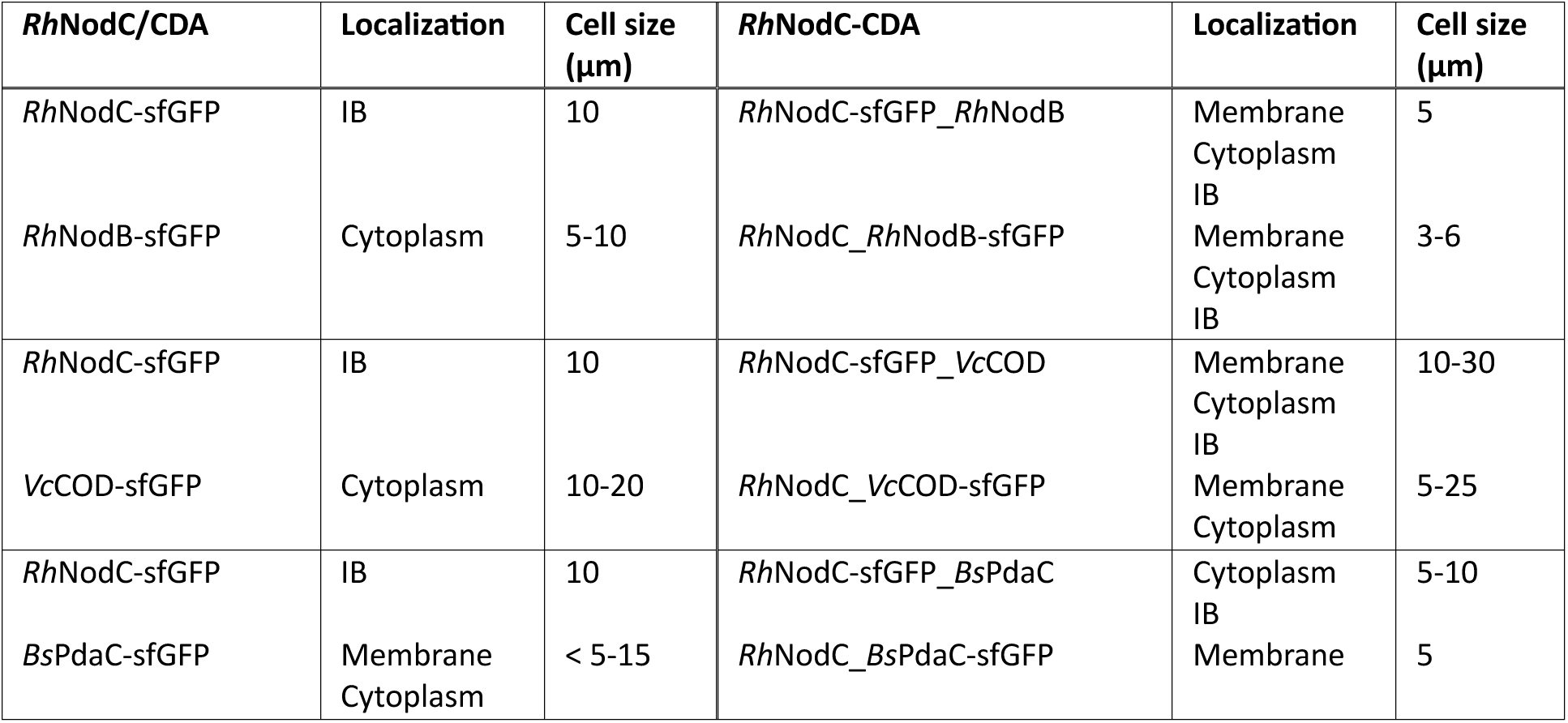
Details of observations made during confocal light scanning microscopy of RhNodC and the different CDAs. On the left-hand side, characteristics of RhNodC or the CDAs expressed on their own can be found, while the right-hand side represents characteristics when RhNodC is co-expressed with the CDAs. The accompanying microscopy images can be found in Figure 5 and Supplementary Figures 2-3. More details on the strains and plasmids are given in Table 4 and Supplementary Table 8. CDA = chitin deacetylase; RhNodC = chitin synthase from Rhizobium sp. GRH2; RhNodB = chitin deacetylase from Rhizobium sp. GRH2; VcCOD = chitin deacetylase from Vibrio cholerae; BsPdaC = chitin deacetylase from Bacillus subtilis; sfGFP = superfolder green fluorescent protein; IB = Inclusion bodies.

| **RhNodC/CDA** | **Localization** | **Cell size ($\mu\text{m}$)** | **RhNodC-CDA** | **Localization** | **Cell size ($\mu\text{m}$)** |
| --- | --- | --- | --- | --- | --- |
| *RhNodC*-sfGFP | IB | 10 | *RhNodC*-sfGFP_*RhNodB* | Membrane<br>Cytoplasm<br>IB | 5 |
| *RhNodB*-sfGFP | Cytoplasm | 5-10 | *RhNodC*_ *RhNodB*-sfGFP | Membrane<br>Cytoplasm<br>IB | 3-6 |
| *RhNodC*-sfGFP | IB | 10 | *RhNodC*-sfGFP_*VcCOD* | Membrane<br>Cytoplasm<br>IB | 10-30 |
| *VcCOD*-sfGFP | Cytoplasm | 10-20 | *RhNodC*_ *VcCOD*-sfGFP | Membrane<br>Cytoplasm | 5-25 |
| *RhNodC*-sfGFP | IB | 10 | *RhNodC*-sfGFP_*BsPdaC* | Cytoplasm<br>IB | 5-10 |
| *BsPdaC*-sfGFP | Membrane<br>Cytoplasm | < 5-15 | *RhNodC*_ *BsPdaC*-sfGFP | Membrane | 5 |

**Table 4:** Bacterial strains used in this study. E. coli = Escherichia coli; KO = knockout.

| Strain | Genetic Background | Purpose | Source |
| --- | --- | --- | --- |
| Top10 | Invitrogen™ One Shot™ Top10<br>Chemically Competent *E. coli* | General cloning | Invitrogen |
| 3KO | *E. coli* K12 MG1655<br>$\Delta chbBCARFG\Delta chiA\Delta nagZ$ | Expression strain | In house |

Finally, *E. coli* cells usually have a length of around 2 µm and a width of 0.5 µm. However, the microscopy images revealed that all cells are elongated (Table 3), which indicates that the strains are under stress [76,77]. The cells in this study had a length between 5 – 10 µm and strains expressing *Vc*COD reached a length of up to 30 µm (see Table 3, Figure 5 and supplementary Figures 2-3). Filamentation occurs when cell growth and DNA replication continue, yet cell division is inhibited [78]. Stress responses connected to this phenotype are the membrane stress response (σ^E^), SOS response (DNA damage), general stress response (σ^S^) and stringent response [47]. The latter two responses were tested in the stress promoter assay but showed no significant activation (Supplementary Figure 1). The exact causes and (stress) mechanisms related to filamentation are still not fully understood, so possibly other stresses might be linked to this phenomenon, or the aforementioned stresses are activated post-transcriptionally and thus not detected in the stress promoter assay.

*Rh*NodC and all of the CDAs located either in the membrane or in IBs when co-expressed, matching the results of the stress promoter assay (Figure 4). For the membrane stress response (*cpx*P), all strains had an increased mKate2 production compared to the control (3KO+*nonCDS*+pInd_*cpx*P). For all but two strains (3KO+*Vc*COD and 3KO+*Rh*NodC_*Bs*Pdac) this increase was significant (One-way ANOVA, p = 1.05E-23; all statistical results can be found in Supplementary Table 7). The results indicate that activation of the *cpx*P promoter is not solely induced by membrane stress, as proteins located in IBs and in the cytoplasm (e.g., *Rh*NodC (IB) and *Rh*NodB (cytoplasm)) also induce the Cpx pathway [45]. The *ibp*A promoter also showed increased response for all strains except for 3KO+*Rh*NodC_*Rh*NodB, even though microscopy (Figure 5) showed formation of IBs for both *Rh*NodC and *Rh*NodB (One-way ANOVA, p = 2.31E-29, all p-values adjusted for multiple comparison by Tukey HSD can be found in Supplementary Table 7). However, membrane stress (*cpx*P) was detected. Since the proteins localize at different positions within the cell, IBs may not be the primary site of stress. Consequently, the activation of *ibp*A could be limited, or the turnover of mKate2 might have been insufficient for detection. Again, proteins located in the cytoplasm (e.g., *Vc*COD, p = 2.02E-13) or membrane (e.g., *Bs*PdaC/*Rh*NodB, increased but not significant, p*_Bs_*_PdaC_ = 7.05E-01/p*_Rh_*_NodB_ = 2.50E-01) also had increased mKate2 production for *ibp*A, as the heat shock response is connected to several other stress responses in the cell [47].

For the first time, we successfully achieved *in vivo* production of partially acetylated chitooligosaccharides (paCOS) through the co-expression of three, different chitin deacetylates (*Rh*NodB, *Vc*COD, and *Bs*PdaC) with *Rh*NodC in *E. coli*. This represents a significant step toward expanding the current COS portfolio. However, co-expression presented several challenges, including reduced total (pa)COS production, halted production after the exponential phase, and incomplete conversion of fully acetylated A5 to paCOS. Production of additional paCOS was also observed in a CDA-dependent manner. To systematically investigate the underlying bottlenecks, we conducted an extensive characterization of the underlying issues by integrating a stress promoter analysis and confocal microscopy. The stress promoter analysis revealed activation of both *ibp*A and *cpx*P stress responses, linked to IB formation and membrane stress, respectively. Notably, we employed a diverse set of stress-responsive promoters covering a broad metabolic stress landscape, demonstrating the utility of this tool for metabolic engineering and host optimization. By identifying specific cellular stress responses, this approach enables a more targeted strategy for strain improvement. Confocal microscopy further provided insights into cellular stress, revealing cell elongation and protein co-localization, suggesting potential *Rh*NodC-CDA interactions or CDA-A5 interactions affecting functionality. Moving forward, efforts should focus on strategies to alleviate these stress-related bottlenecks through metabolic engineering or dynamic pathway regulation. Addressing these bottlenecks will be crucial for improving the controlled *in vivo* production of (pa)COS, ultimately advancing its industrial applications.

## 3. Materials and Methods

### 3.1. Chemicals and molecular biology

Enzymes and related products are purchased from New England Biolabs (County Road, Ipswich, MA, USA), other products from Sigma-Aldrich (Brusselsesteenweg, Overijse, Belgium) unless stated differently. Standard molecular biology methods were applied as described by Sambrook et al. [79].

### 3.2. Strains and media

Bacterial strains used in this study are given in Table 4. For the construction and storage of plasmids, One Shot^TM^ Top10 Chemically Competent *E. coli* (Invitrogen, Carlsbad, California, USA) was used. For the different assays, a previously developed *E. coli* K12 MG1655 Δ*chb*BCARFGΔ*chi*AΔ*nag*Z (3KO) strain was used, where the three knockouts were made using the method developed by Datsenko and Wanner [80]. This strain was transformed with the corresponding plasmids (Supplementary Table 8) via electroporation or heat shock.

Lysogenic broth (LB), consisting of 10 g/L Tryptone (BioKar Diagnostics, Allonne, France), 5 g/L Yeast Extract (Becton Dickinson, Erembodegem-Dorp, Erembodegem, Belgium) and 5 g/L NaCl was used to grow the strains during cloning and transformation processes, or for the precultures for microtiter plate (MTP) and shake flask experiments. For LB-agar plates, 12 g/L agar (BioKar Diagnostics, Allonne, France) was added. If selection was necessary, antibiotics were added to LB(-agar) in 1000x dilution, with the following stock solutions of the respective antibiotics: 50 mg/mL kanamycin, 25 mg/mL chloramphenicol and 100 mg/mL ampicillin. Cultures were grown at 30°C and liquid cultures were shaken at 200 rpm (5 cm orbit, LS-X, Adolf Kühner AG, Switzerland).

For MTP and shake flask experiments, strains were grown on minimal medium with glucose (MMGlc) (composition given in Table 5). To avoid Maillard reactions during autoclaving, the carbon source and magnesium sulphate (20% v/v of total solution) were separately filter sterilized (250 mL filter unit, 0.2 μm PES membrane; VWR International BVBA, Researchpark Haasrode, Geldenaaksebaan, Leuven, Belgium). The other salts and components (remaining 80%) were set to a pH of 7 using KOH before being autoclaved (121°C, 30 min, 1.02 atm). The two solutions were mixed and supplemented with the filter sterilized (Sterile syringe filter 0.2 μm PES; VWR) vitamin mix and molybdate solution (Table 5).

**Table 5:** Left. Medium composition of minimal medium glucose used for microtiter plate and shake flask experiments. **Right.** Composition of the vitamin mix and molybdate solution added to the minimal medium.

| Compound | Amount | Compound | Amount |
| --- | --- | --- | --- |
| NH<sub>4</sub>Cl | 2.00 g/L | **Vitamin mix** |  |
| (NH<sub>4</sub>)SO<sub>4</sub> | 5.00 g/L | Na<sub>2</sub>EDTA.2H<sub>2</sub>O | 0.4 g/L |
| KH<sub>2</sub>PO<sub>4</sub> | 2.993 g/L | H<sub>3</sub>BO<sub>3</sub> | 0.0311 g/L |
| K<sub>2</sub>HPO<sub>4</sub> | 7.315 g/L | thiamine.HCl | 1.01 g/L |
| MOPS | 8.372 g/L | ZnCl<sub>2</sub> | 0.94 g/L |
| NaCl | 0.5 g/L | CoCl<sub>2</sub>.6H<sub>2</sub>O | 0.5 g/L |
|  | Dissolve in 800 mL/L dH<sub>2</sub>O | CuCl<sub>2</sub>.2H<sub>2</sub>O | 0.38 g/L |
| Glucose.H<sub>2</sub>O | 16.5 g/L | MnCl<sub>2</sub>.4H<sub>2</sub>O | 1.59 g/L |
| MgSO<sub>4</sub>.7H<sub>2</sub>O | 0,5 g/L | CaCl<sub>2</sub> | 3.77 g/L |
|  | Dissolve in 200 mL/L dH<sub>2</sub>O | FeCl<sub>2</sub>.4H<sub>2</sub>O | 3.6 g/L |
| Vitamin mix | 1 mL/L | **Molybdate** |  |
| Molybdate | 100 µL/L | Na<sub>2</sub>MoO<sub>4</sub>.2H<sub>2</sub>O | 0.967 g/L |
|  |  | SeO<sub>2</sub> | 3 g/L |

### 3.3. Plasmid construction

An overview of all the plasmids used in this study is given in Supplementary Table 8. Plasmids for the production and stress promoter assay that contain *RhNod*C and/or the CDAs and the plasmids to study localization of the proteins consist of a medium copy pBR322 origin of replication (ori) and a kanamycin resistance gene (the sequences of the plasmid backbones can be found in Supplementary Tables 9 and 10). The plasmids were constructed using Circular Polymerase Extension Cloning (CPEC) [81], with the Q5 High-Fidelity DNA polymerase (Bioké, Leiden, The Netherlands). The indicator plasmids were assembled in the MoBioS system [82]. Stress promoters were ordered as linear fragments from Integrated DNA Technologies BVBA (Interleuvenlaan, Leuven Belgium or GeneArt (Thermo Fisher Scientific). Assembly into the backbone (the sequences of the plasmid backbones can be found in Supplementary Tables 9 and 10) with the low copy pSC101 ori and chloramphenicol resistance gene was done with Golden Gate (GG) based assembly using the PaqCI restriction enzyme.

All polymerase chain reactions (PCR) performed were done using either PrimeStar HS (Takara, Westburg, Leusden, The Netherlands) or PrimeStar Gxl DNA polymerase (Takara). DNA fragments were purified using the innuPREP PCRpure Kit (Analytic Jena AG, Jena, Germany).

*E. coli* Top10 was transformed with the newly constructed plasmids and the presence of the plasmids in the colonies was verified by colony PCR. To check the DNA sequences of the newly constructed plasmids, the positive colonies were grown overnight (30°C, 200 rpm) and prepped using the QIAprep Spin Miniprep Kit (Qiagen, Venlo, The Netherlands). Purified plasmids were sent to Macrogen (Macrogen Inc., Amsterdam, The Netherlands) for Sanger sequencing. After verification, the plasmids were transformed into the correct genetic background.

### 3.4. *In vivo* fluorescence measurements

For the stress promoter assay, the 3KO strain (see Table 4) was transformed with the plasmids containing *RhNod*C and the different CDAs, together with the different indicator plasmids (Supplementary Table 8). 3KO containing two similar plasmids with a non-coding sequence (3KO+*nonCDS*+pInd junk) was taken as a control for background fluorescence of *E. coli*. 3KO expressing an empty plasmid (3KO+*nonCDS*) together with the indicator plasmids served as a benchmark for the activation of the stress promoters in unstressed cells.

For the stress promoter assay, precultures from five single colonies per strain were grown in 150 μL LB in a Transparent CELLSTAR^©^ polystyrene 96-well microtiter plates, covered with a clear polystyrene lid with condensation rings (Greiner Bio-One Vilvoorde, Belgium) for 24 h at 30°C and 800 rpm on the Compact Digital Microplate Shaker (Thermo Fisher Scientific, Erembodegem, Belgium). Subsequently, precultures were diluted 300x into 150 μL MMGlc in a Black μCLEAR^©^ polystyrene 96-well microtiter plate covered with a clear polystyrene lid with condensation rings (Greiner Bio-One Vilvoorde, Belgium). Plates were then transferred into the Tecan M200 Pro (MNano^+^) plate reader (Tecan Benelux, Mechelen, Belgium) (30°C, 200 rpm). Optical density at 600 nm (OD_600_) was followed up to quantify growth, while for detection of fluorescence, an excitation and emission wavelength of 588 nm and 633 nm for mKate2 and 480 nm and 510 nm for sfGFP were applied.

### 3.5. Partially acetylated chitooligosaccharide production assay

To analyze (pa)COS production, 3 biological replicates (single colonies) per strain were grown on LB for 24 h at 30°C and 200 rpm (5 cm orbit, LS-X, Adolf Kühner AG, Switzerland). Next, they were inoculated (1% v/v) in 50 mL MMGlc with antibiotics (250 mL shake flask). To follow up growth, samples for OD_600_ (200 μL - 1 mL, diluted to be in the linear range OD_600_ = 0 - 1) were regularly taken and measured with the Fisherbrand^TM^ Cell Density Meter (Thermo Fisher Scientific, Erembodegem, Belgium). In the exponential, late exponential and stationary phase (pa)COS samples were taken. For this, 1.5 mL of culture was transferred to a 2.0 mL tube and centrifuged for 10 min at 20 627 x g (Sigma 1-16, Sigma Laborzentrifugen GmbH; Osterode am Harz, Germany). The pellet was dried and then stored at-80°C until further use. To prepare the samples for quantification, 150 μL of 60 % acetonitrile (AcN) (Chem-Lab NV, Industriezone “De Arend” 2, Zedelgem, Belgium) in milli-Q water (mQ) was added to the pellet and vortexed until completely dissolved. The samples were centrifuged for 10 min at 20 627 x g (Sigma 1-16, Sigma Laborzentrifugen GmbH). Finally, 100 μL of the supernatant was transferred to a vial (VerexTM Vial, 9 mm Screw Top, μVial i3 (Qsert), Clear 33, No Patch; Phenomex) and closed off with a cap (VerexTM Cap (pre-assembled), 9 mm, Screw top, w/ PTFE/Silicone presplit septa, blue; Phenomex) for analysis on the Shimadzu High-performance liquid chromatography (HPLC) system (Shimadzu Prominence-i LC 2030C Plus, Shimadzu, Jette, Belgium). A chitinpentaose standard was purchased from Neogen (Ireland) while a monodeacetylated chitopentaose (DP = 5, DA = 20%) standard was obtained from the European Nano3Bio project (European 7^th^ Framework program, GA613931). Details for the HPLC run can be found in Table 6. Separation was done by hydrophilic interaction chromatography (HILIC), with the Kinetix 2.6 μm HILIC 100A column (2.6 μm, 4.6 mm x 150 mm) (Phenomenex, Utrecht, The Netherlands). The pre-column consists of the following two components: SecurityGuard ULTRA Cartridges, UHPLC HILIC 4.6mm ID Columns - AJ0-8772 (Phenomenex) and SecurityGuard ULTRA Holder, for UHPLC Columns 2.1 to 4.6mm ID - AJ0-9000 (Phenomenex).

**Table 6:** High-performance liquid chromatography (HPLC) analysis for the quantification of chitooligosaccharide production. Characteristics of the HPLC and its method are given. AcN = acetonitrile; FA = formate; A5 = fully acetylated chitopentaose; paCOS (DP=5, DDA=20%) = chitopentaose with a degree of deacetylation of 20%; column T = column temperature; ELSD = evaporative light scattering detector.

| General characteristics |  | Method |  |  |
| --- | --- | --- | --- | --- |
| **Column** | Kinetix 2.6 μm HILIC 100A column<br>(2.6 μm, 4.6 mm x 150 mm) | **Time (min)** | **%A** | **%B** |
| **Elution type** | Gradient | 0-8 | 85 | 15 |
| **Eluent A** | 100% AcN + 0.1 v/v % FA | 8-20 | 55 | 45 |
| **Eluent B** | 10 mM NH<sub>4</sub>-FA + 0.1 v/v % FA | 20-27 | 50 | 50 |
| **Flow** | 1.0 mL/min | 27-30 | 85 | 15 |
| **Column T** | 45°C | 30-40 | 85 | 15 |
| **Method length** | 40 min | **Component** | **Elution time (min)** |  |
| **Detection** | ELSD | A5 | 15.25 - 16.50 |  |
| **Injection volume** | 2 μL | paCOS<br>(DP=5,<br>DDA=20%) | 17.25 - 18.75 | |

### 3.6. Confocal light scanning microscopy

Strains for microscopy were grown overnight on LB (30°C, 200 rpm) (5 cm orbit, LS-X, Adolf Kühner AG, Switzerland) and were then sub cultivated into MMGlc and grown for another 24 h (30°C, 200 rpm, 5 cm orbit). Cultures were diluted 10x in phosphate-buffered saline and 500 μL of this dilution was dyed with 1 μL of red FM4-64 membrane dye (red) (Thermo Fisher Scientific, Erembodegem, Belgium) to be able to locate the cells. Cultures were then applied on a μ-slide 8 well coverslip (ibidi, Martinsried, Germany) coated with a 0.01 % poly-L-lysine solution and incubated for 30 min at room temperature and pipetted dry thereafter. Subsequently, a few droplets of mounting medium (1 % n-propyl-gallate in glycerol) were added to each well. Samples were then analyzed using the Zeiss LSM 780 confocal scanning light microscopy with Airyscan technology, a service provided by the Bio-imaging facility at VIB Ghent. Both the red (membrane) and green (proteins) channels were visualized. Microscopy images were analyzed using Fiji (an image processing package of ImageJ). To analyze the localization of the proteins, an intensity plot of the cross-section of single cells was made.

### 3.7. Data and statistical analysis

Data analysis was performed in Python using the pandas package (https://pandas.pydata.org/) unless stated differently.

To determine the maximal growth rate (μ_max_), OD_600_ values were fitted to the Richards growth model (Equation 1.1) [83]. In this equation, y is the log-transformed OD_600_ values in function of the time t (h). A is the maximal value of y, ν a shape parameter, μ_max_ the maximal growth rate (h^-1^) and λ represents the lag time (h). Fitting was done with the scipy.optimize.curve_fit function in Python and as such μ can be determined. The geometric average and standard error are shown in the plots.

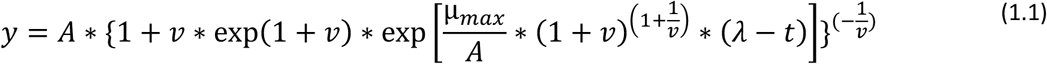

For peak integration of the chromatograms, the LabSolutions software (Shimadzu, Jette, Belgium) was used. COS titers were determined based on a calibration curve for A5 and paCOS (DP=5, DDA=20%) (0.1 - 0.25 - 0.5 - 0.75 - 1.0 (-1.5 - 2.0) g/L)). COS production (g/L) was calculated based on the area of the peaks, the calibration curve and the concentration of the sample. COS composition (%) was determined solely based on the areas of the peaks. Error bars represent the standard error of the mean.

Fluorescence measurements for the stress promoter assay were corrected for background fluorescence and OD_600_ of the medium and the cell culture (Equation 1.2). MMGlc without a cell culture was used to correct for background fluorescence of the medium (FP and OD, respectively) and was averaged over all time points and replicates. An *E. coli* strain (3KO+*nonCDS*) that does not express any fluorescence genes was used to correct for background fluorescence of *E. coli* (FP_Blank_ and OD_Blank_, respectively) and was averaged per time point for all replicates.

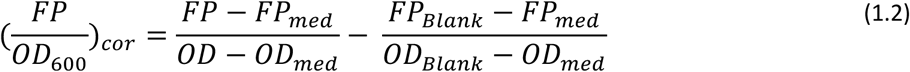

In the plots, the weighted mean of the corrected fluorescence is given. Corrected fluorescence in the stationary phase was determined over a time span of 21 data points around entry into the stationary phase. Error propagation rules were applied to calculate the standard error of the mean.

For statistical analysis, a significance level of 0.05 was applied and the scipy.stats package in Python was used. Pairwise comparison of strains was done using a two-sided two-sample T-test (scipy.stats.ttest_ind). If more than two samples had to be compared, one-way ANOVA was done (stats.scipy.f)_oneway) and consequently Tukey HSD to correct for multiple comparison (statsmodels.stats.multicomp.pairwise_tukeyhsd). Because the geometrical mean of μ_max_ is used, these values were log-transformed before conducting statistical analysis.

## 4. Author contributions

S. Snoeck, C. Guidi and M. De Mey contributed to the conceptualization of the manuscript. H. Malik, S. Snoeck, W. Lips and D. Duchi collected and analyzed the data. S. Snoeck and H. Malik wrote the manuscript that was critically reviewed by W. Demeester, C. Guidi and M. De Mey.

## 6. Acknowledgments

S.S. holds a PhD grant [Grant no. 11H5122N] from FWO (Fonds Wetenschappelijk Onderzoek – Vlaanderen). This research was also supported by a Catalisti VLAIO-ICON project’Encaps2Control’ (HBC.2019.0122), FWO Research project’SynSysBio4COS’ (G0B8118), FWO Bioeconomy Research Project’MyCOS’ (GOG3422N) and BOF IOP’Immunokeys’ (BOF/IOP/2022/069). Confocal scanning light microscopy, with Airyscan technology was provided by the Bio-imaging Facility at VIB Ghent. We would like to thank the UGent Core Facility ‘HTS for Synthetic Biology’ for training, support and access to the instrument park.

## 7. Appendix A. Supplementary data

The supplementary data file contains:

‒ Supplementary tables 1-12
‒ Supplementary figures 1-4

