## Supplementary data for "Understanding bottlenecks in the microbial production of partially acetylated chitooligosaccharides"

**Supplementary Table 1:** List of DNA sequences used in this study (coding sequences, promoter, 5’UTR and terminator sequences). All sequences are given from 5’ to 3’. The core promoter (-35 box to TSS) is given together with the upstream context of the promoter, the -35 and -10 box are underlined. RBS = ribosome binding site; UTR = untranslated region; TSS = transcription start site; Rh = *Rhizobium* sp. GRH2, Vc = *Vibrio cholerae*, Bs = *Bacillus subtilis*.

| **Gene** | **Coding sequence** | |
|---|---|---|
| *Rhnod*C | 1 | ATG GAC CTG CTG AAC ACG ATT GGT ATT GGT GCT GTC TCC TGC TAC GCT CTG CTG TCA ACG GCT CAT AAG TCG ATG CAA ACC CTG |
| | 85 | TAT GCC CAG CCG AAA GAT CAA AGC TCT GCA TCA GAA GAC TTT GCT TTC CTG CCG TCG GTG GAT ATT ATC GTT CCG TGT TAT AAC |
| | 169 | GAA AAT CCG CAT ACC TTT AGC GAA TGC CTG GCG TCT ATT GCC AAC CAG GAT TAT GCG GGC AAA CTG CGT GTG TAC GTG GTT GAT |
| | 253 | GAC GGT AGT GCC AAT CGT GAA AAG CTG GAA CGC GTT CAT CAC ACC TAC GCA GGC GAT CCG CGT TTT GAC TTC ATC CTG CTG CGT |
| | 337 | GAA AAC GTG GGT AAG CGT AAG GCA CAG ATT GCA GCA ATC CGT GGC AGT TCC GGT GAT CTG GTG CTG AAT GTT GAT AGC GAC TCT |
| | 421 | ACC CTG GCA TCA GAC GTC GTG ACG AAA CTG GCT CTG AAG ATG CAG AAC CCG GAA ATT GGC GCA GCT ATG GGT CAA CTG ACC GCG |
| | 505 | TCT AAC CGT AAT GAT ACC TGG CTG ACG CGC CTG ATC GAC ATG GAA TAT TGG CTG GCC TGT AAT GAA GAA CGT GCA GCA CAG GCA |
| | 589 | CGT TTT GGT GCA GTG ATG TGC TGT TGC GGT CCG TGC GCA ATG TAT CGT CGC TCA GCT CTG CTG TCG CTG CTG GAT CAG TAC GAA |
| | 673 | AGC CAA TTT TTC CGT GGC AAA CCG TCT GAT TTT GGT GAA GAC CGC CAT CTG ACC ATT CTG ATG CTG AAG GCG GGC TTC CGT ACG |
| | 757 | GAT TAT GTT CCG GAC GCC ATC GCA GCT ACC GTT GTC CCG GAT CGT ATG GGT CCG TAC CTG CGC CAG CAA CTG CGT TGG GCA CGC |
| | 841 | AGC ACC TTC CGT GAT ACG CTG CTG GCT CTG CGT CTG CTG CCG GGT CTG GAT CAC TAT ATT ACG CTG GAC GTT ATC GGT CAG AAC |
| | 925 | CTG GGT CCG CTG CTG CTG GCA CTG GCT GTC CTG ACC GGT GTC CTG CAA GTG GCA CTG ACC GCT ACG GTC CCG CTG TGG ACC GTG |
| | 1009 | ATG ATG ATT GCA TCA ATG ACG ATG ATC CGT TGT GCA GTT GCA GCA GTC CGT GCA CGT CAG CTG CGC TTT CTG GTT TTC TCG CTG |
| | 1093 | CAT ACC CCG ATT AAC CTG TTT TTC CTG CTG CCG ATG AAA GCG TAC GCC CTG TGC ACG CTG AGT AAC TCC GAT TGG CTG AGT CGC |
| | 1177 | TCA TCG CCG GCG AAT AAA ACC TCC GCC GGC GGT GAA CAC CCG ACC ACG GAA GCA AGT GCT GGC GGT ACC TCC GGC AAC GCG ACG |
| | 1261 | CCG CTG CGT CGC CTG AAC CTG GCT CGT GAC TCC TCT ACC GTT ACC CCG GCT GGT GTC TAC TCC GAT GAT TGA |
| *Rhnod*B | 1 | ATG ACG CAC TTT GAC TGC CTG AGC CGT GTC CAA TCG ACC TGT AGC AAC TCG ACG GGT GGC CGC AAT GTG TAT CTG ACC TTT GAC |
| | 85 | GAC GGT CCG AAC CCG CTG AGT ACC CCG GAA ATT CTG GAT GTG CTG GAA AAA CAT CGT GTT CCG GCA ACG TTT TTC GTC ATC GGC |
| | 169 | GCT TAT GCG GCC GAA CAG CCG CAA CTG ATT CGT CGC ATG ATC GTT GAA GGT CAC GAA GTC GCG AAC CAT ACC ATG ACG CAC CCG |
| | 253 | GAT CTG AGT CGT TGC GGC CCG TCC GAC GTT CGC CAT GAA ATT CTG GAA GCG AAT CGC GTG GTT ACC ATG GCA TGT CCG CAG GCT |
| | 337 | TCC ATT CGT CAC GTG CGC GCC CCG TAC GGT ATC TGG ACG GAT GAA GTC CTG ACC ACG AGC GCC GGT GTG GGT CTG GCA CCG CTG |
| | 421 | CAT TGG AGC GTT GAT CCG CGT GAC TGG TCG CGT CCG GGT GTC GAT GCA ATT GTC GAC ACC GTG CTG AGC AGC GTG CAG CCG GGT |
| | 505 | GCT ATC GTT CTG CTG CAC GAT GGT TGC CCG CCG GAC GAA GTT CGT CCG GGT ACC GAA GAT AGC CTG CGC GAC CAA ACC GTG ACG |
| | 589 | GCG CTG TCT CGT CTG ATC CCG GCC CTG CAT GAA CGT GGC TTC GGT ATC TGT GCT CTG CCG CAA CAT CCG CTG ACC AAA GAG CT |

**Supplementary Table 1:** Continued (1).

| Gene | Coding sequence |
|---|---|
| *VcCOD* | 1 ATG AAC AGC ACG CCG AAA GGC ACG ATC TAT CTG ACG TTT GAT GAT GGC CCG GTG AAC GCC TCG GTG GAA GTG ATT AAA GTC CTG<br>85 AAC CAG GGC GGT GTT AAA GCC ACC TTT TAT TTC AAC GCA TGG CAT CTG GAT GGC ATT GGT GAC GAA AAT GAA GAT CGT GCA CTG<br>169 GAA GCT CTG AAA CTG GCA CTG GAC AGT GGC CAT ATT GTG GGT AAC CAC TCC TAC GAT CAC ATG ATC CAC AAT TGC GTT GAA GAA<br>253 TTT GGC CCG ACC AGC GGT GCG GAT TGT AAC GCC ACG GGT AAT CAT CAG ATC CAC TCA TAC CAA GAC CCG GTG CGT GAT GCG GCC<br>337 TCG TTC GAA CAG AAC CTG ATT ACC CTG GAA AAA TAT CTG CCG ACG ATC CGC AGC TAT CCG AAC TAC AAA GGC TAT GAA CTG GCG<br>421 CGT CTG CCG TAC ACC AAT GGT TGG CGC GTC ACG AAA CAT TTT CAG GCA GAC GGC CTG TGC GCT ACC TCA GAT AAT CTG AAA CCG<br>505 TGG GAA CCG GGT TAT GTG TGT GAT CCG GCT AAC CCG TCA AAT TCG GTG AAA GCG TCG ATT CAG GTT CAA AAC ATC CTG GCC AAT<br>589 CAG GGC TAC CAA ACC CAC GGT TGG GAT GTT GAC TGG GCT CCG GAA AAC TGG GGC ATT CCG ATG CCG GCG AAT AGC CTG ACC GAA<br>673 GCG GTC CCG TTT CTG GCC TAT GTG GAC AAA GCA CTG AAC AGC TGC TCT CCG ACC ACG ATT GAA CCG ATC AAT TCT AAA ACC CAG<br>757 GAA TTC CCG TGT GGT ACG CCG CTG CAT GCG GAT AAA GTC ATC GTG CTG ACC CAC GAC TTT CTG TTC GAA GAT GGC AAA CGT GGC<br>841 ATG GGT GCC ACG CAA AAC CTG CCG AAA CTG GCA GAA TTT ATT CGC ATC GCG AAA GAA GCC GGT TAC GTG TTC GAT ACC ATG GAC<br>925 AAT TAT ACG CCG CGT TGG AGT GTG GGC AAA ACC TAC CAG GCC GGT GAA TAC GTT CTG TAT CAA GGC GTG GTT TAT AAA GCA GTC<br>1009 ATT TCC CAC ACG GCT CAG CAA GAC TGG GCG CCG AGC AGC ACC AGT TCC CTG TGG ACG AAC GCA GAT CCG GCT ACC AAC TGG ACG<br>1093 CTG AAT GTT AGC TAT GAA CAG GGC GAT ATC GTC AAT TAC AAA GGT AAA CGC TAT CTG GTT TCT GTC CCG CAT GTT TCG CAG CAA<br>1177 GAC TGG ACG CCG GAC ACC CAA AAT ACC CTG TTC ACC GCA CTG G |
| *Bspda*C | 1 ATG GAA GAA ACA GTT GAT CCG AAT CAG AAA GTT ATT GCC CTG ACC TTT GAT GAT GGT CCG AAT CCG GCA ACC ACC AAT CAG<br>82 ATT CTG GAT AGT CTG AAA AAA TAC AAA GGC CAC GCC ACC TTT TTT GTT CTG GGT AGC CGT GTT CAG TAT TAT CCG GAA ACC<br>163 CTG ATT CGT ATG CTG AAA GAG GGT AAC GAA GTT GGT AAT CAT AGC TGG TCA CAT CCG CTG CTG ACC CGT CTG AGC GTT AAA<br>244 GAA GCC CTG AAA CAA ATT AAT GAT ACC CAG GAC ATC ATC GAG AAA ATT AGC GGT TAT CGT CCG ACC CTG GTT CGT CCG CCT<br>325 TAT GGT GGT ATT AAT GAT GAA CTG CGT AGC CAG ATG AAA ATG GAT GTT GCA CTG TGG GAT GTG GAT CCG GAA GAT TGG AAA<br>406 GAT CGT AAC AAA AAA ACC ATT GTG GAT CGC GTT ATG AAT CAA GCC GGT GAT GGT CGT ACC ATT CTG ATT CAT GAT ATT TAT<br>487 CGT ACC AGC GCA GAT GCA GCC GAT GAA ATT ATC AAA AAA CTG ACC GAT CAG GGT TAT CAG CTG GTT ACC GTT AGC CAG CTG<br>568 GAA GAG GTT AAA AAA CAG CGT GAA GCA AAA |

**Supplementary Table 1:** Continued (2).

| **Promoter** | **DNA sequence** |
|---|---|
| $P_{14}$ p[BR322] | GCTTTACATCGCTTCAGTGCTTGTACCCATCTGATGCACGCCATCGGAACGCCTTCATTCTATAAGTTTCTTGACATCTTGGCCGGCATATGGTA<br>TAATAGGGCC |
| $P_{22}$ p[BR322] | CAATCCCAAAATGGCGAAGTCCTTAGCGGTAATCCCAGCGTCGTTGTTCTGGGACCACTCGAAGAAGATGTTGACATTTTGGAATAGATGTGA<br>TATAATGTGTAC |
| $P_{14}$ p[15A] | AACAGATAAAACGAAAGGCCCAGTCTTTCGACTGAGCCTTTCGTTTTATTTGATGCCTTAATTAAAAGGGGTCTCGACTCAGGAACTCTTCATTCT<br>ATAAGTTTCTTGACATCTTGGCCGGCATATGGTATAATAGGGCC |
| **Stress promoter**<br>**Promoter + 5'UTR** | **DNA sequence**<br>-35 and -10 box(es) underlined as reported in literature |
| $P_{junk}$ | AAATGAACGATTTCTTAGTCGGCGTTATAGTAAGTCACTCTTTTTCAGCGGTATTTTAAAGATGAGAAAGCGATGGTCAAGCGTGGTCTGCCTGAA<br>GTCT |
| $P_{\mathit{rel}\mathrm{A}}$ | GCTGATGATTTTGCGCCATACCGCACCGCTAAGTTCGGCAGATCGCGAAAAACTGGAACGCTTTTCGCATTCTGAAGGCCTGGATCTGTATCTCGCC<br>CCCGATAGTGAGATACTCGAAACCGTCTCTGGTGAGATGCCCTGGTATGACTCAAACGGGTTGCGCTTAACTTTTAGCCCGCGCGATTTTATTCAGG<br>TCAATGCGGGTGTGAACCAAAAAATGGTAGCGCGTGCGTTGGAATGGCTGGATGTGCAACCTGAAGATCGCGTACTGGATCTGTTCTGCGGTATG<br>GGCAACTTTACACTGCCATTGGCGACACAAGCTGCCAGTGTGGTCGGTGTAGAAGGTGTTCCGGCGCTGGTGGAAAAAGGCCAGCAGAATGCGC<br>GTCTTAATGGCTTACAGAATGTGACGTTTTATCACGAAAATCTTGAAGAAGATGTCACAAAGCAGCCGTGGGCGAAAAACGGCTTCGATAAAGTG<br>TTGCTGGACCCGGCGCGAGCAGGTGCCGCAGGTGTTATGCAGCAAATTATAAAACTGGAACCTATTCGTATAGTTTATGTATCCTGTAACCCTGCA<br>ACGCTGGCTCGGGATAGCGAAGCGTTATTAAAAGCAGGATATACCATTGCGCGACTGGCGATGCTGGATATGTTCCCACACACGGGACATCTGGA<br>ATCGATGGTACTTTTCTCGCGCGTTAAATAGTTGCGA TTTGCCGATTTCGGCAGGTCTGGTCCCTAAAGGAGAGGACG |
| $P_{spoT}$ | CTTCGATACCGCGTTGACCGATTTGAAGACCATTATTCGCGCCGAACGTCTGCGCATGAGCCGCCAAAAGCAGCGTCATGACGCTTTAATCAGCAA<br>ATTGTTGGCAGACTGAACCTGATTTCAGTATCATGCCCAGTCATTTCTTCACCTGTGGAGCTTTTTAAGT |
| $P_{\mathit{suh}\mathrm{B}}$ | GCTGCCAGGGCAATCGCCTGGGAGTCGGGTTTCACCAGTGGATTAACCAGCCACAGATTGGTTAATCCCATTGTTTTCATGGCACGGGCAACAGAA<br>CCCATATTGCCGGTGTGTGACGTCTCCACCAGCACAATTCGAATATTTTGCAGCATTGTCTTTCTTCATCTAAAGATTATTCACGCATCTTATCATAAA<br>ACGAAGACAGATGCCGATCTCGCTGCTATACTCTGCGCCGTTTTCCCGTTCTTTAACATCCAGTGAGAGAG ACCG |
| $P_{\mathit{ira}\mathrm{P}}$ | CTTAAAAACCTATCCCGTCTAACACAAAGTGCATACATTACCACGACAAAACGGGGGATTCGCGGCCTTCTGAAAGATTGTTGCAATCTTCTGCTGA<br>CAAAGCGTGCAACGTACTGGTGAAGAAAGTGCGTTATCTCAAAGATGTGCGCAAGATCACAAAAATGATGAACGGGAAGCTAATTTATTCCTGGC<br>TTAAATGGCCATGCGGTGAGTTTTTTTCTCTTAATTATAAGTTAACGAAGAGAATATATTTCATAACTTTTATTTATAATAAAGGTTGATAATTAAAA<br>GCCTATATTTTGTGTGGGTAATTATTTAAATAAGAGAAACGTTTCGCTGGTAATCAAACAAAAAATATTTGCGCAAAGTATTTCCTTTGTCATAAAAA<br>TAATACTTCCAGACACTATGAAGTTGTGAAACATAATGTTAACTTCTCCATACTTT GGATAAGGAAATACAGAC |
| $P_{\mathit{dsr}\mathrm{A}}$<br>+ 5'UTR *pro*B | CACATTTCTATTCATAAGTAGCGTTAATCATTCATATGGCGAATATTTTCTTGTCAGCGAAAAAAATTGCGGATAAGGTGATGAAGGGAGACCACA<br>ACGGTTTCCCTCTACAAATAATTTTGTTTAACTTTTACTAGAGTCACACAGGAAAGTACTAG |

**Supplementary Table 1:** Continued (3).

| **Stress promoter**<br>**Promoter + 5'UTR** | **DNA sequence**<br>-35 and -10 box(es) underlined as reported in literature |
|---|---|
| $P_{rpoS}$ | CCTTGCTCAGCGCAACAATATTCAGGCACCATACGCGCTGAACGTTGGTCAGACCTTGCAGGTGGGTAATGCTTCCGGTACGCCAATCACTGGCG<br>GAAATGCCATTACCCAGGCCGACGCAGCAGAGCAAGGAGTTGTGATCAAGCCTGCACAAAATTCCACCGTTGCTGTTGCGTCGCAACCGACAATT<br>ACGTATTCTGAGTCTTCGGGTGAACAGAGTGCTAACAAAATGTTGCCGAACAACAAGCCAACTGCGACCACGGTCACAGCGCCTGTAACGGTACC<br>AACAGCAAGCACAACCGAGCCGACTGTCAGCAGTACATCAACCAGTACGCCTATCTCCACCTGGCGCTGGCCGACTGAGGGCAAAGTGATCGAA<br>ACCTTTGGCGCTTCTGAGGGGGGCAACAAGGGGATTGATATCGCAGGCAGCAAAGGACAGGCAATTATCGCGACCGCAGATGGCCGCGTTGTTT<br>ATGCTGGTAACGCGCTGCGCGGCTACGGTAATCTGATTATCATCAAACATAATGATGATTACCTGAGTGCCTACGCCCATAACGACACAATGCTGG<br>TCCGGGAACAACAAGAAGTTAAGGCGGGGCAAAAAATAGCGACCATGGGTAGCACCGGAACCAGTTCAACACGCTTGCATTTTGAAATTCGTTA<br>CAAGGGGAAATCCGTAAACCCGCTGCGTTATTTGCCGCAGCGATAAATGGCGGAACCAGGCTTTTGCTTGAATGTTCCGTCAAGGGATCACGGGT<br>AGGAGCCACCTT |
| $P_{ibpA}$ | CAGTCTATGCAATAGACCATAAACTGCAAAAAAAAGTCCGCTGATAAGGCTTGAAAAGTTCATTTCCAGACCCATTTTTACATCGTAGCCGATGAG<br>GACGCGCCTGATGGGTGTTCTGGCTACCTGACCTGTCCATTGTGGAAGGTCTTACATTCTCGCTGATTTCAGGAGCTAT TGATT |
| $P_{rpoH}$ | TTGTCATCGGCGGTTGCGGAAGTGGCACAGGTTTTCGGAACGAAGTTTGATATCAATGGCTTATCATTCGATGAATGCCTGCTATTGCTGCTGGTA<br>TGCTCGATGATTGGCTGGGTGGCAGCGTGGCTTGCCACGGTACAACATTTACGCCACTTTACGCCTGAATAATAAAAGCGTGTTATACTCTTTCCC<br>TGCAATGGGTTCCGTAGCAGGGAAAGAGACCCCGTTGTCTCTTCCCGGTATTTCATCTCTATGTCACATTTTGTGCGTAATTTATTCACAAGCTTGC<br>ATTGAACTTGTGGATAAAATCACGGTCTGATAAAACAGTGAATGATAACCTCGTTGCTCTTAAGCTCTGGCACAGTTGTTGCTACCACTGAAGCGC<br>CAGAAGATATCGATTGAGAGGATTTGA |
| $P_{cpxP}$ | AAATACCTCCGAGGCAGAAATTACGTCATCAGACGTCGCTAATCCATGACTTTACGTTGTTTTACACCCCCTGACGCATGTTTGCAGCCTGAATCGT<br>AAACTCTCTATCGTTGAATCGCGACAGAAAGATTTTGGGAGCAAATG |

| **5'UTR** | **DNA sequence** |
|---|---|
| P14 5'UTR<br>p[BR322] | CCTCTAGAAATAATTTTGTTTAACTTTAAGAAGGAGATATACATA |
| P14 5'UTR<br>p[15A] | CCTCTAGAAATAAGTTTTGTTTAACTTTAAGAAGGAGATATACATA |
| P22 5'UTR<br>RBS(oNodB5) | ATATCCAACAAACCATAGAGTAGGACGTCTCTTT |
| P22 5'UTR<br>RBS(oVc6) | ATATCCAAACGAAATCCACCCCGTTAAGAAAGGCGGTATAGT |

**Supplementary Table 1:** Continued (4).

| **5'UTR** | **DNA sequence** |
|---|---|
| P22 5'UTR RBS(syn) | ATATCCAGGTATAATAAGTAAAATGGGTTATCG |
| P22 5'UTR RBS(BsaIRBS_TIR321) | CGTCACACTACCCGCTAGCGACGTACCAGGTCCAAGGAGGTCTCT |
| BCD16 | GGGCCCAAGTTCACTTAAAAAGGAGATCAACAATGAAAGCAATTTTCGTACTGAAACATCTTAATCATGCTTAGGAGTCTTTCT |
| **Terminator** | |
| TT5-T7term | ACAACCCTCAAGAGAAAATGTAATCACACTGGCTCACCTTCGGGTGGGCCTTTCTGCGTTTATAAGGAGACACTTTATGTTTAAGAAG |
| TT7-M13centralT | AAAGCAAGCTGATAAACCGATACAATTAAAGGCTCCTTTTGGAGCCTTTTTTTTTGGAGATTTTCAACATGAAAAAATTATTATT |

**Supplementary Table 2:** Results of the statistical analysis for the comparison of the (partially) acetylated chitooligosaccharide production ($g*L^{-1}$) between different strains. One-way ANOVA was performed and Tukey HSD was conducted to correct for multiple comparison. Significant values ($p < 0.05$) are indicated in bold. More details on the strains and plasmids can be found in Table 4 and Supplementary Table 8. *Rh*NodC = chitin oligosaccharide synthase from *Rhizobium* sp. GRH2; *Rh*NodB = chitin deacetylase from *Rhizobium* sp. GRH2; *Vc*COD = chitin deacetylase from *Vibrio cholerae*; *Bs*PdaC = chitin deacetylase from *Bacillus subtilis*; A5 = fully acetylated chitin pentaose; paCOS (DP=5, DDA=20%) = chitin pentaose with a degree of deacetylation of 20 %.

| | | **Exponential** | | **Late exponential** | | **Stationary** | |
|---|---|---|---|---|---|---|---|
| | | **A5** | **paCOS (DP=5, DDA=20%)** | **A5** | **paCOS (DP=5, DDA=20%)** | **A5** | **paCOS (DP=5, DDA=20%)** |
| **ANOVA** | **F-value** | 365.70 | 42.53 | 1355.12 | 47.39 | 560.00 | 149.48 |
| | **p-value** | **1.12E-12** | **3.18E-07** | **4.47E-16** | **1.73E-07** | **1.04E-11** | **7.13E-09** |
| **Strain 1** | **Strain 2** | | | | | | |
| *Rh*NodC | *Rh*NodC_ *Bs*PdaC | **2.63E-12** | 1.00E+00 | **5.55E-16** | **2.60E-05** | **5.05E-11** | **1.37E-06** |
| *Rh*NodC | *Rh*NodC_ *Vc*COD | **3.98E-08** | **4.80E-04** | **5.33E-15** | **2.15E-06** | **1.56E-10** | **6.35E-08** |
| *Rh*NodC | *Rh*NodC_ *Rh*NodB | **6.02E-09** | **1.46E-06** | **3.33E-15** | **6.81E-06** | **1.28E-10** | **1.29E-07** |
| *Rh*NodC | *nonCDS* | **2.63E-12** | 1.00E+00 | **0.00E+00** | 1.00E+00 | **8.31E-12** | 1.00E+00 |
| *Rh*NodC_ *Bs*PdaC | *Rh*NodC_ *Vc*COD | **2.40E-09** | **4.80E-04** | **3.29E-04** | 2.91E-01 | **2.02E-02** | **5.10E-03** |
| *Rh*NodC_ *Bs*PdaC | *Rh*NodC_ *Rh*NodB | **1.38E-08** | **1.46E-06** | **4.43E-03** | 8.51E-01 | 5.59E-02 | **3.30E-02** |
| *Rh*NodC_ *Bs*PdaC | *nonCDS* | 1.00E+00 | 1.00E+00 | **7.36E-06** | **2.60E-05** | **2.24E-04** | **1.37E-06** |
| *Rh*NodC_ *Vc*COD | *Rh*NodC_ *Rh*NodB | 1.43E-01 | **4.54E-03** | 5.62E-01 | 8.79E-01 | 9.61E-01 | 7.32E-01 |
| *Rh*NodC_ *Vc*COD | *nonCDS* | **2.40E-09** | **4.80E-04** | **2.26E-08** | **2.15E-06** | **4.87E-06** | **6.35E-08** |
| *Rh*NodC_ *Rh*NodB | *nonCDS* | **1.38E-08** | **1.46E-06** | **8.07E-08** | **6.81E-06** | **8.56E-06** | **1.29E-07** |

**Supplementary Table 3:** Results of the statistical analysis for the comparison of the specific (partially) acetylated chitooligosaccharide production ($g*L^{-1}*OD_{600}^{-1}$) between different strains. One-way ANOVA was performed and Tukey HSD was conducted to correct for multiple comparison. Significant values ($p < 0.05$) are indicated in bold. More details on the strains and plasmids can be found in Table 4 and Supplementary Table 8. RhNodC = chitin oligosaccharide synthase from *Rhizobium* sp. GRH2; RhNodB = chitin deacetylase from *Rhizobium* sp. GRH2; VcCOD = chitin deacetylase from *Vibrio cholerae*; BsPdaC = chitin deacetylase from *Bacillus subtilis*; A5 = fully acetylated chitin pentaose; paCOS (DP=5, DDA=20%) = chitin pentaose with a degree of deacetylation of 20%; $OD_{600}$ = optical density measured at 600 nm.

| | | Exponential | | Late exponential | | Stationary | |
|---|---|---|---|---|---|---|---|
| | | **A5** | **paCOS (DP=5, DDA=20%)** | **A5** | **paCOS (DP=5, DDA=20%)** | **A5** | **paCOS (DP=5, DDA=20%)** |
| **ANOVA** | **F-value** | 134.33 | 29.43 | 378.27 | 33.98 | 382.59 | 238.69 |
| | **p-value** | **4.19E-10** | **2.44E-06** | **9.13E-13** | **1.11E-06** | **6.88E-11** | **7.12E-10** |
| **Strain 1** | **Strain 2** | | | | | | |
| *Rh*NodC | *Rh*NodC_ *Bs*PdaC | **1.37E-08** | 1.00E+00 | **2.91E-12** | **9.22E-03** | **4.37E-10** | **2.92E-06** |
| *Rh*NodC | *Rh*NodC_ *Vc*COD | **6.76E-05** | **1.26E-02** | **1.48E-11** | **4.10E-04** | **2.11E-09** | **1.09E-07** |
| *Rh*NodC | *Rh*NodC_ *Rh*NodB | **3.30E-05** | **3.91E-05** | **5.04E-11** | **2.37E-05** | **4.19E-08** | **1.68E-09** |
| *Rh*NodC | *nonCDS* | **1.37E-08** | 1.00E+00 | **6.30E-13** | 1.00E+00 | **4.49E-11** | 1.00E+00 |
| *Rh*NodC_ *Bs*PdaC | *Rh*NodC_ *Vc*COD | **1.22E-05** | **1.26E-02** | **8.68E-03** | 4.05E-01 | **1.10E-02** | **4.91E-03** |
| *Rh*NodC_ *Bs*PdaC | *Rh*NodC_ *Rh*NodB | **2.38E-05** | **3.91E-05** | **1.03E-04** | **1.38E-02** | **6.11E-06** | **6.85E-07** |
| *Rh*NodC_ *Bs*PdaC | *nonCDS* | 1.00E+00 | 1.00E+00 | **5.19E-03** | **9.22E-03** | **1.57E-04** | **2.92E-06** |
| *Rh*NodC_ *Vc*COD | *Rh*NodC_ *Rh*NodB | 9.92E-01 | **1.99E-02** | 1.00E-01 | 3.29E-01 | **5.02E-04** | **3.83E-05** |
| *Rh*NodC_ *Vc*COD | *nonCDS* | **1.22E-05** | **1.26E-02** | **1.13E-05** | **4.10E-04** | **2.73E-06** | **1.09E-07** |
| *Rh*NodC_ *Rh*NodB | *nonCDS* | **2.38E-05** | **3.91E-05** | **5.52E-07** | **2.37E-05** | **4.07E-08** | **1.68E-09** |

**Supplementary Table 4:** Results of the statistical analysis for the comparison of the (partially) acetylated chitooligosaccharide productivity ($g*L^{-1}*h^{-1}$) between different strains. One-way ANOVA was performed and Tukey HSD was conducted to correct for multiple comparison. Significant values ($p < 0.05$) are indicated in bold. More details on the strains and plasmids can be found in Table 4 and Supplementary Table 8. RhNodC = chitin oligosaccharide synthase from *Rhizobium* sp. GRH2; RhNodB = chitin deacetylase from *Rhizobium* sp. GRH2; VcCOD = chitin deacetylase from *Vibrio cholerae*; BsPdaC = chitin deacetylase from *Bacillus subtilis*; A5 = fully acetylated chitin pentaose; paCOS (DP=5, DDA=20%) = chitin pentaose with a degree of deacetylation of 20%.

| | | **Exponential** | | **Late exponential** | | **Stationary** | |
|---|---|---|---|---|---|---|---|
| | | **A5** | **paCOS (DP=5, DDA=20%)** | **A5** | **paCOS (DP=5, DDA=20%)** | **A5** | **paCOS (DP=5, DDA=20%)** |
| **ANOVA** | **F-value** | 1011.97 | 73.17 | 4336.19 | 147.51 | 734.81 | 172.24 |
| | **p-value** | **2.56E-15** | **1.44E-08** | **4.20E-19** | **2.42E-10** | **2.68E-12** | **3.55E-09** |
| **Strain 1** | **Strain 2** | | | | | | |
| *Rh*NodC | *Rh*NodC_ *Bs*PdaC | **4.11E-15** | 1.00E+00 | **0.00E+00** | **1.57E-08** | **1.62E-11** | **2.55E-07** |
| *Rh*NodC | *Rh*NodC_ *Vc*COD | **9.65E-11** | **1.74E-07** | **0.00E+00** | **1.02E-09** | **5.00E-11** | **1.13E-08** |
| *Rh*NodC | *Rh*NodC_ *Rh*NodB | **1.16E-13** | **1.36E-06** | **0.00E+00** | **9.82E-06** | **1.22E-11** | **3.55E-06** |
| *Rh*NodC | *nonCDS* | **4.11E-15** | 1.00E+00 | **0.00E+00** | 1.00E+00 | **2.65E-12** | 1.00E+00 |
| *Rh*NodC_ *Bs*PdaC | *Rh*NodC_ *Vc*COD | **5.41E-12** | **1.74E-07** | **1.36E-06** | **8.91E-03** | **9.88E-03** | **1.40E-03** |
| *Rh*NodC_ *Bs*PdaC | *Rh*NodC_ *Rh*NodB | **2.05E-07** | **1.36E-06** | **1.23E-02** | **1.12E-04** | 7.70E-01 | **2.75E-02** |
| *Rh*NodC_ *Bs*PdaC | *nonCDS* | 1.00E+00 | 1.00E+00 | **1.92E-08** | **1.57E-08** | **8.40E-05** | **2.55E-07** |
| *Rh*NodC_ *Vc*COD | *Rh*NodC_ *Rh*NodB | **2.40E-09** | 2.82E-01 | **3.77E-08** | **7.98E-07** | **1.86E-03** | **2.15E-05** |
| *Rh*NodC_ *Vc*COD | *nonCDS* | **5.41E-12** | **1.74E-07** | **4.36E-11** | **1.02E-09** | **1.67E-06** | **1.13E-08** |
| *Rh*NodC_ *Rh*NodB | *nonCDS* | **2.05E-07** | **1.36E-06** | **5.64E-07** | **9.82E-06** | **3.16E-04** | **3.55E-06** |

**Supplementary Table 5:** Results of the statistical analysis for the comparison of the total chitooligosaccharide production (g*$L^{-1}$) (A5 and paCOS (DP=5, DDA=20%) ) between different strains. One-way ANOVA was performed and Tukey HSD was conducted to correct for multiple comparison. Significant values ($p < 0.05$) are indicated in bold. More details on thestrains and plasmids can be found in Table 4 and Supplementary Table 8. RhNodC = chitin oligosaccharide synthase from *Rhizobium* sp. GRH2; RhNodB = chitin deacetylase from *Rhizobium* sp. GRH2; VcCOD = chitin deacetylase from *Vibrio cholerae*; BsPdaC = chitin deacetylase from *Bacillus subtilis*; A5 = fully acetylated chitin pentaose; paCOS (DP=5, DDA=20%) = chitin pentaose with a degree of deacetylation of 20%.

| | | **Exponential** | **Late exponential** | **Stationary** |
|---|---|---|---|---|
| **ANOVA** | **F-value** | 73.04 | 207.14 | 275.36 |
| | **p-value** | **1.46E-08** | **3.26E-11** | **3.51E-10** |
| **Strain 1** | **Strain 2** | | | |
| *Rh*NodC | *Rh*NodC_ *Bs*PdaC | **4.42E-06** | **5.97E-10** | **8.56E-09** |
| *Rh*NodC | *Rh*NodC_ *Vc*COD | 9.80E-01 | **7.20E-09** | **1.19E-07** |
| *Rh*NodC | *Rh*NodC_ *Rh*NodB | **2.39E-02** | **2.94E-09** | **6.51E-08** |
| *Rh*NodC | *nonCDS* | **4.42E-06** | **8.67E-12** | **1.39E-10** |
| *Rh*NodC_ *Bs*PdaC | *Rh*NodC_ *Vc*COD | **2.15E-06** | **1.19E-02** | **3.36E-03** |
| *Rh*NodC_ *Bs*PdaC | *Rh*NodC_ *Rh*NodB | **1.25E-07** | 1.26E-01 | **1.59E-02** |
| *Rh*NodC_ *Bs*PdaC | *nonCDS* | 1.00E+00 | **7.78E-06** | **4.58E-06** |
| *Rh*NodC_ *Vc*COD | *Rh*NodC_ *Rh*NodB | 7.58E-02 | 7.18E-01 | 8.25E-01 |
| *Rh*NodC_ *Vc*COD | *nonCDS* | **2.15E-06** | **1.36E-07** | **1.30E-07** |
| *Rh*NodC_ *Rh*NodB | *nonCDS* | **1.25E-07** | **4.65E-07** | **2.47E-07** |

**Supplementary Table 6:** Results of the statistical analysis for the comparison of the chitooligosaccharide production (g*L$^{-1}$) between different growth phases. One-way ANOVA was performed and Tukey HSD was conducted to correctfor multiple comparison. Significant values ($p < 0.05$) are indicated in bold. More details on the strains and plasmids can be found in Table 4 and Supplementary Table 9. RhNodC = chitin oligosaccharide synthase from *Rhizobium* sp. GRH2; *Rh*NodB = chitin deacetylase from *Rhizobium* sp. GRH2; *Vc*COD = chitin deacetylase from *Vibrio cholerae*; *Bs*PdaC = chitin deacetylase from *Bacillus subtilis*; E = exponential phase; L = late exponential phase; S = stationary phase; A5 = fully acetylated chitin pentaose; paCOS (DP=5, DDA=20%) = chitin pentaose with a degree of deacetylation of 20%; nan = not a number (no statistical comparison can be made because no chitooligosaccharides were detected).

| | | ***nonCDS*** | | ***Rh*NodC** | | ***Rh*NodC_ *Rh*NodB** | |
|---|---|---|---|---|---|---|---|
| | | A5 | paCOS (DP=5, DDA=20%) | A5 | paCOS (DP=5, DDA=20%) | A5 | paCOS (DP=5, DDA=20%) |
| **ANOVA** | **F-value** | nan | nan | 603.98 | nan | 2.88 | 3.45 |
| | **p-value** | nan | nan | **1.21E-07** | nan | 1.33E-01 | 1.01E-01 |
| **Growth phase 1** | **Growth phase 2** | | | | | | |
| E | L | nan | nan | **1.03E-07** | nan | 1.29E-01 | 2.97E-01 |
| E | S | nan | nan | **1.32E-06** | nan | 2.80E-01 | 1.46E-01 |
| L | S | nan | nan | **5.20E-05** | nan | 8.16E-01 | 8.41E-01 |

| | | ***Rh*NodC_ *Vc*COD** | | ***Rh*NodC_ *Bs*PdaC** | |
|---|---|---|---|---|---|
| | | A5 | paCOS (DP=5, DDA=20%) | A5 | paCOS (DP=5, DDA=20%) |
| **ANOVA** | **F-value** | 68.61 | 133.14 | 3172.68 | 634.34 |
| | **p-value** | **7.35E-05** | **1.07E-05** | **8.43E-10** | **1.04E-07** |
| **Growth phase 1** | **Growth phase 2** | | | | |
| E | L | **5.77E-05** | **1.64E-03** | **1.41E-09** | **1.08E-07** |
| E | S | **2.58E-03** | 5.23E-02 | **1.72E-09** | **5.02E-07** |
| L | S | **2.72E-03** | **3.40E-02** | 1.39E-01 | **6.01E-04** |

**Supplementary Table 7:** Statistical one-way ANOVA analysis and Tukey HSD for the activation of stress promoters *ibp*A and *cpx*P. Significant values ($p < 0.05$) are indicated in bold. More details on the strains and plasmids can be found in Table 4 and Supplementary Table 8. RhNodC = chitin oligosaccharide synthase from *Rhizobium* sp. GRH2; *Rh*NodB = chitin deacetylase from *Rhizobium* sp. GRH2; *Vc*COD = chitin deacetylase from *Vibrio cholerae*; *Bs*PdaC =chitin deacetylase from *Bacillus subtilis*.

| | | ***ibp*A** | ***cpx*P** |
|---|---|---|---|
| **ANOVA** | **F-value** | 190.21 | 90.11 |
| | **p-value** | **2.31E-29** | **1.05E-23** |
| **Strain 1** | **Strain 2** | | |
| *Bs*PdaC | *nonCDS* | 7.05E-01 | **0.00E+00** |
| *Vc*COD | *nonCDS* | **2.02E-13** | 1.00E+00 |
| *Rh*NodB | *nonCDS* | 2.50E-01 | **5.00E-15** |
| *Rh*NodC | *nonCDS* | **3.28E-14** | **1.43E-04** |
| *Rh*NodC_*Bs*PdaC | *nonCDS* | 6.23E-01 | 1.00E+00 |
| *Rh*NodC_*Vc*COD | *nonCDS* | **2.58E-04** | **1.46E-02** |
| *Rh*NodC_*Rh*NodB | *nonCDS* | 1.00E+00 | **3.27E-11** |
| *Bs*PdaC | *Vc*COD | **2.54E-11** | **0.00E+00** |
| *Bs*PdaC | *Rh*NodB | 9.99E-01 | 9.45E-02 |
| *Bs*PdaC | *Rh*NodC | **2.82E-12** | **2.32E-12** |
| *Bs*PdaC | *Rh*NodC_*Bs*PdaC | 1.00E+00 | **0.00E+00** |
| *Bs*PdaC | *Rh*NodC_*Vc*COD | **4.92E-02** | **3.89E-14** |
| *Bs*PdaC | *Rh*NodC_*Rh*NodB | 6.55E-01 | **6.87E-06** |
| *VcCOD* | *Rh*NodB | **2.15E-10** | **9.55E-15** |
| *VcCOD* | *Rh*NodC | 9.98E-01 | **2.83E-04** |
| *VcCOD* | *Rh*NodC_*Bs*PdaC | **3.61E-11** | 1.00E+00 |
| *VcCOD* | *Rh*NodC_*Vc*COD | **5.27E-07** | **2.61E-02** |
| *Rh*NodC_*Vc*COD | *Rh*NodC_*Rh*NodB | **2.03E-04** | **3.78E-06** |

| | | ***ibp*A** | ***cpx*P** |
|---|---|---|---|
| **ANOVA** | **F-value** | 190.21 | 90.11 |
| | **p-value** | **2.31E-29** | **1.05E-23** |
| **Strain 1** | **Strain 2** | | |
| *VcCOD* | *Rh*NodC_*Rh*NodB | **1.67E-13** | **6.08E-11** |
| *Rh*NodB | *Rhnod*C | **2.21E-11** | **1.78E-08** |
| *Rh*NodB | *Rh*NodC_ *Bs*PdaC | 1.00E+00 | **2.75E-14** |
| *Rh*NodB | *Rh*NodC_*Vc*COD | 2.53E-01 | **1.70E-10** |
| *Rh*NodB | *Rh*NodC_*Rh*NodB | 2.16E-01 | 6.37E-02 |
| *Rh*NodC | *Rh*NodC_ *Bs*PdaC | **3.95E-12** | **9.56E-04** |
| *Rh*NodC | *Rh*NodC_*Vc*COD | **4.26E-08** | 8.63E-01 |
| *Rh*NodC | *Rh*NodC_*Rh*NodB | **2.82E-14** | **5.13E-04** |
| *Rh*NodC_ *Bs*PdaC | *Rh*NodC_*Vc*COD | 6.68E-02 | 6.97E-02 |
| *Rh*NodC_ *Bs*PdaC | *Rh*NodC_*Rh*NodB | 5.72E-01 | **1.90E-10** |

**Supplementary Table 8: Plasmids used in this study.** The coding sequences, promoter, 5' untranslated region and terminator sequences can be found in the Supplementary Table 1. Sequences of the backbones can be found in Supplementary Tables 10-12. ori = origin of replication; AB = antibiotics resistance; Prom = promoter; RBS = ribosome binding site; CDS = coding sequence; Term = terminator; Kan = kanamycin; Chl = chloramphenicol; Rh = Rhizobium sp. GRH2; Vc = Vibrio cholerae; Bs = Bacillus subtilis; BCD = bicistronic design.

| **Plasmid** | **Plasmid details: p[ori][AB][Prom-RBS-CDS-Term]$_n$** |
|---|---|
| **Production plasmids** | |
| *nonCDS* | p[BR322][Kan][*nonCDS*] |
| *Rhnod*C | p[BR322][Kan][P14-RBS(T7)-*Rhnod*C-T(TT5-T7term)][*nonCDS*] |
| *Rhnod*C_*Rhnod*B | p[BR322][Kan][P14-RBS(T7)-*Rhnod*C-T(TT5-T7term)][P22-RBS(oNodB5)-*Rhnod*B-T(TT7-M13centralT)] |
| *Rhnod*C_*VcCOD* | p[BR322][Kan][P14-RBS(T7)-*Rhnod*C-T(TT5-T7term)][P22-RBS(oVc6)-*VcCOD*-T(TT7-M13centralT)] |
| *Rhnod*C_*Bspda*C | p[BR322][Kan][P14-RBS(T7)-*Rhnod*C-T(TT5-T7term)][P22-RBS(syn)-*Bspda*C-T(TT7-M13centralT)] |
| **Stress promoters** | |
| *nonCDS* | p[BR322][Kan][*nonCDS*] |
| *Rhnod*C | p[BR322][Kan][P14-RBS(T7)-*Rhnod*C-T(TT5-T7term)][*nonCDS*] |
| *Rhnod*C_*RhnodB* | p[BR322][Kan][P14-RBS(T7)-*Rhnod*C-T(TT5-T7term)][P22-RBS(oNodB5)-*Rhnod*B-T(TT7-M13centralT)] |
| *Rhnod*C_*VcCOD* | p[BR322][Kan][P14-RBS(T7)-*Rhnod*C-T(TT5-T7term)][P22-RBS(oVc6)-*VcCOD*-T(TT7-M13centralT)] |
| *Rhnod*C_*Bspda*C | p[BR322][Kan][P14-RBS(T7)-*Rhnod*C-T(TT5-T7term)][P22-RBS(syn)-*Bpda*C-T(TT7-M13centralT)] |
| *Rhnod*B | p[BR322][Kan][P22-RBS(oNodB5)-*Rhnod*B-T(TT7-M13centralT)] |
| *VcCOD* | p[BR322][Kan][P22-RBS(oVc6)-*VcCOD*-T(TT7-M13centralT)] |
| *Bspda*C | p[BR322][Kan][P22-RBS(syn)-*Bpda*C-TT7-M13centralT] |
| pInd_junk | p[SC101][Chl][PPTFBS junk -RBS(junk)-mKate-T(FAB391)]-[P22 -RBS(BsaIRBS TIR321)-TF junk(L3)-T(TT3-rrnD1-T1)] |
| pInd_*rel*A | p[SC101][Chl][P*rel*A-RBS(*rel*A)-*mKate*-T(FAB391)]-[P22-RBS(BsaIRBS TIR321)-TF junk(L3)-T(TT3-rrnD1-T1)] |
| pInd_*spo*T | p[SC101][Chl][P*spo*T-RBS(*spo*T)-*mKate*-T(FAB391)]-[P22-RBS(BsaIRBS TIR321)-TF junk(L3)-T(TT3-rrnD1-T1)] |
| pInd_*suh*B | p[SC101][Chl][P*suh*B-RBS(*suh*B)-*mKate*-T(FAB391)]-[P22-RBS(BsaIRBS TIR321)-TF junk(L3)-T(TT3-rrnD1-T1)] |
| pInd_*ira*P | p[SC101][Chl][P*ira*P-RBS(*ira*P)-*mKate*-T(FAB391)]-[P22-RBS(BsaIRBS TIR321)-TF junk(L3)-T(TT3-rrnD1-T1)] |
| pInd_*dsr*A | p[SC101][Chl][P*dsr*A-RBS(*dsr*A)-*mKate*-T(FAB391)]-[P22-RBS(BsaIRBS TIR321)-TF junk(L3)-T(TT3-rrnD1-T1)] |
| pInd_*rpo*S | p[SC101][Chl][P*rpo*S-RBS(*rpo*S)-*mKate*-T(FAB391)]-[P22-RBS(BsaIRBS TIR321)-TF junk(L3)-T(TT3-rrnD1-T1)] |
| pind_*ibp*A | p[SC101][Chl][P*ibp*A-RBS(*ibp*A)-*mKate*-T(FAB391)]-[P22-RBS(BsaIRBS TIR321)-TF junk(L3)-T(TT3-rrnD1-T1)] |
| pInd_*rpo*H | p[SC101][Chl][P*rpo*H-RBS(*rpo*H)-*mKate*-T(FAB391)]-[P22-RBS(BsaIRBS TIR321)-TF junk(L3)-T(TT3-rrnD1-T1)] |
| pInd_*cpx*P | p[SC101][Chl][P*cpx*P-RBS(*cpx*P)-*mKate*-T(FAB391)]-[P22-RBS(BsaIRBS TIR321)-TF junk(L3)-T(TT3-rrnD1-T1)] |

**Supplementary Table 8: Plasmids used in this study.** The coding sequences, promoter, 5' untranslated region and terminator sequences can be found in the Supplementary Table 1. Sequences of the backbones can be found in Supplementary Tables 10-12. ori = origin of replication; AB = antibiotics resistance; Prom = promoter; RBS = ribosome binding site; CDS = coding sequence; Term = terminator; Kan = kanamycin; Chl = chloramphenicol; Rh = Rhizobium sp. GRH2; Vc = Vibrio cholerae; Bs = Bacillus subtilis; BCD = bicistronic design. Continued (1)

| **Plasmid** | **Plasmid details: p[ori][AB][Prom-RBS-CDS-Term]$_n$** |
|---|---|
| **Localization** | |
| *sfGFP* | p[BR322][Kan][P14-RBS(T7)-*sfGFP*-His-T(TT5-T7term)] |
| *RhnodC-sfGFP* | p[BR322][Kan][P14-RBS(T7)-*Rhnod*C-*sfGFP*-His-T(TT5-T7term)] |
| *RhnodB-sfGFP* | p[BR322][Kan][P22-RBS(oNodB5)-*Rhnod*B-*sfGFP*-His-T(TT7-M13centralT)] |
| *VcCOD-sfGFP* | p[BR322][Kan][P22-RBS(oVc6)-*VcCOD-sfGFP*-His-T(TT7-M13centralT)] |
| *BspdaC-sfGFP* | p[BR322][Kan][P22-RBS(syn)-*Bspda*C-*sfGFP*-His-T(TT7-M13centralT)] |
| *RhnodC_RhnodB-sfGFP* | p[BR322][Kan][P14-RBS(T7)-*Rhnod*C-T(TT5-T7term)][P22-RBS(oNodB5)-*Rhnod*B-*sfGFP*-His-T(TT7-M13centralT)] |
| *RhnodC_VcCOD-sfGFP* | p[BR322][Kan][P14-RBS(T7)-*Rhnod*C-T(TT5-T7term)][P22-RBS(oVc6)-*VcCOD-sfGFP*-His-T(TT7-M13centralT)] |
| *RhnodC_BspdaC-sfGFP* | p[BR322][Kan][P14-RBS(T7)-*Rhnod*C-T(TT5-T7term)][P22-RBS(syn)-*Bspda*C-*sfGFP*-His-T(TT7-M13centralT)] |
| *RhnodC-sfGFP_RhnodB* | p[BR322][Kan][P14-RBS(T7)-*Rhnod*C-*sfGFP*-His-T(TT5-T7term)][P22-RBS(oNodB5)-*Rhnod*B-T(TT7-M13centralT)] |
| *RhnodC-sfGFP_VcCOD* | p[BR322][Kan][P14-RBS(T7)-*Rhnod*C-*sfGFP*-His-T(TT5-T7term)][P22-RBS(oVc6)-*VcCOD*-T(TT7-M13centralT)] |
| *RhnodC-sfGFP_BspdaC* | p[BR322][Kan][P14-RBS(T7)-*Rhnod*C-*sfGFP*-His-T(TT5-T7term)][P22-RBS(syn)-*Bspda*C-T(TT7-M13centralT)] |

**Supplementary Table 9: Plasmid backbone p[BR322]with kanamycin resistance used throughout this study.** The sequences of interest are inserted at NNNNNNNNNN and are surrounded by a non-coding sequence. The sequences of the regulatory elements and gene sequences can be found in Supplementary Table 1.

| Plasmid | Sequence |
|---|---|
| **p[BR322][Kan]** | 1<br>AGTCCTAGGATGCTAGCTATGTGGGCTTACATGGCGATAGCTAGACTGGGCGGTTTTATGGACA<br>GCAAGC<br>71<br>GAACCGGAATTGCCAGCTGGGGCGCCCTCTGGTAAGGTTGGGAAGCCCTGCAAAGTAAACTGGA<br>TGGCTT<br>141<br>TCTTGCCGCCAAGGATCTGATGGCGCAGGGGATCAAGATCTGATCAAGAGACAGGATGAGGAT<br>CGTTTCG<br>211<br>CATGATTGAACAAGATGGATTGCACGCAGGTTCTCCGGCCGCTTGGGTGGAGAGGCTATTCGGC<br>TATGAC<br>281<br>TGGGCACAACAGACAATCGGCTGCTCTGATGCCGCCGTGTTCCGGCTGTCAGCGCAGGGGCGCC<br>CGGTTC<br>351<br>TTTTTGTCAAGACCGACCTGTCCGGTGCCCTGAATGAACTGCAGGACGAGGCAGCGCGGCTATC<br>GTGGCT<br>421<br>GGCCACGACGGGCGTTCCTTGCGCAGCTGTGCTCGACGTTGTCACTGAAGCGGGAAGGGACTGG<br>CTGCTA<br>491<br>TTGGGCGAAGTGCCGGGGCAGGATCTCCTGTCATCTCACCTTGCTCCTGCCGAGAAAGTATCCA<br>TCATGG<br>561<br>CTGATGCAATGCGGCGGCTGCATACGCTTGATCCGGCTACCTGCCCATTCGACCACCAAGCGAA<br>ACATCG<br>631<br>CATCGAGCGAGCACGTACTCGGATGGAAGCCGGTCTTGTCGATCAGGATGATCTGGACGAAGAG<br>CATCAG<br>701<br>GGGCTCGCGCCAGCCGAACTGTTCGCCAGGCTCAAGGCGCGCATGCCCGACGGCGAGGATCTCG<br>TCGTGA<br>771<br>CCCATGCGATGCCTGCTTGCCGAATATCATGGTGGAAAATGGCCGCTTTTCTGGATTCATCGAC<br>TGTGGC<br>841<br>CGGCTGGGTGTGGCGGACCGCTATCAGGACATAGCGTTGGCTACCCGTGATATTGCTGAAGAGC<br>TTGGCG<br>911<br>GCGAATGGGCTGACCGCTTCCTCGTGCTTTACGGTATCGCCGCTCCCGATTCGCAGCGCATCGCC<br>TTCTT<br>981<br>ATCGCCTTCTTGACGAGTTCTTCTGAGCGGGACTCTGGGGTTCGAAATGACCGACCAAGCGACG<br>CCCAAC<br>1051<br>CTGCCATCACGAGATTTCGATTCCACCGCCGCCTTCCCCCCATGAACAGAAATCCCCCTTACACG<br>GAGGC<br>1121<br>ATCAGTGACCAAACAGGAAAAAACCGCCCTTAACATGGCCCGCTTTATCAGAAGCCAGACATTA<br>ACGCTT<br>1191<br>CTGGAGAAACTCAACGAGCTGGACGCGGATGAACAGGCAGACATCTGTGAATCGCTTCACGACC<br>ACGCTG |

| | |
|---|---|
| | 1261<br>ATGAGCTTTACCGCAGCTGCCTCGCGCGTTTCGGTGATGACGGTGAAAACCTCTGACACATGCA<br>GCTCCC<br>1331<br>GGAGACGGTCACAGCTTGCTGTAAGCGGATGCCGGGAGCAGACAAGCCCGTCAGGGCGCGTCAG<br>CGGGTG<br>1401<br>TTGGCGGGTGTCGGGGCGCAGCCATGACCCAGTCACGTAGCGATAGCGGAGTGTATACTGGCTT<br>AACTAT<br>1471<br>GCGGCATCAGAGCAGATTGTACTGAGAGTGCACCATATGCGGTGTGAAATACCGCACAGATGCG<br>TAAGGA<br>1541<br>GAAAATACCGCATCAGGCGCTCTTCCGCTTCCTCGCTCACTGACTCGCTGCGCTCGGTCGTTCGG<br>CTGCG<br>1611<br>GGCGAGCGGTATCAGCTCACTCAAAGGCGGTAATACGGTTATCCACAGAATCAGGGGATAACGC<br>AGGAAA<br>1681<br>GAACATGTGAGCAAAAGGCCAGCAAAAGGCCAGGAACCGTAAAAAGGCCGCGTTGCTGGCGTT<br>TTTCCAT<br>1751<br>AGGCTCCGCCCCCCTGACGAGCATCACAAAAATCGACGCTCAAGTCAGAGGTGGCGAAACCCG<br>ACAGGA<br>1821<br>CTATAAAGATACCAGGCGTTTCCCCCTGGAAGCTCCCTCGTGCGCTCTCCTGTTCCGACCCTGCC<br>GCTTA<br>1891<br>CCGGATACCTGTCCGCCTTTCTCCCTTCGGGAAGCGTGGCGCTTTCTCATAAGCTCACGCTGTAG<br>GTATC<br>1961<br>TCAGTTCGGTGTAGGTCGTTCGCTCCAAGCTGGGCTGTGTGCACGAACCCCCCGTTCAGCCCGA<br>CCGCTG<br>2031<br>CGCCTTATCCGGTAACTATCGTCTTGAGTCCAACCCGGTAAGACACGACTTATCGCCACTGGCA<br>GCAGCC<br>2101<br>ACTGGTAACAGGATTAGCAGAGCGAGGTATGTAGGCGGTGCTACAGAGTTCTTGAAGTGGTGG<br>CCTAACC<br>2171<br>TGGCTACACTAGAAGGACAGTATTTGGTATCTGCGCTCTGCTGAAGCCAGTTACCTTCGGAAAA<br>AGAGTTG<br>2241<br>GTAGCTCTTGATCCGGCAAACAAACCACCGCTGGTAGCGGTGGTTTTTTTGTTTGCAAGCAGCA<br>GATTAC<br>2311<br>GCGCAGAAAAAAAGGATCTCAAGAAGATCCTTTGATCTTTTCTACGGGGTCTGACGCTCAGTGG<br>AACGAA<br>2381<br>AACTCACGTTAAGGGATTTTGGTCATGAGATTATCAAAAAGGATCTTCACCTAGATCCTTTTAA<br>ATTAAA<br>2451<br>AATGAAGTTTTAAATCAATCTAAAGTATATATGAGTAAACTTGGTCTGACAGAGCTGGCACGA<br>CAGGTTT<br><br>2521<br>CCCGACTGGAAATAGACGTCGCCTCAGCTTGTGACGAAAAGTGCCACCTGACGTCATTAATNNN<br>NNNNNN |

**Supplementary Table 10: Plasmid backbone p[SC101] with chloramphenicol resistance used throughout this study.** Backbone of the indicator plasmids used for the stress promoter assay. mKate is indicated in red, the promoters are inserted at NNNNNNNNNN. The stress promoter sequences can be found in Supplementary Table 1.

| Plasmid | Sequence |
|---|---|
| **p[SC101][Chl]** | 1 GTAAGACGGG TAAGCCTGTT GATGATACCG CTGCCTTACT GGGTGCATTA GCCAGTCTGA ATGACCTGTC<br><br>71 ACGGGATAAT CCGAAGTGGT CAGACTGGAA AATCAGAGGG CAGGAACTGC TGAACAGCAA AAAGTCAGAT<br><br>141 AGCACCACAT AGCAGACCCG CCATAAAACG CCCTGAGAAG CCCGTGACGG GCTTTTCTTG TATTATGGGT<br><br>211 AGTTTCCTTG CATGAATCCA TAAAAGGCGC CTGTAGTGCC ATTTACCCCC ATTCACTGCC AGAGCCGTGA<br><br>281 GCGCAGCGAA CTGAATGTCA CGAAAAAGAC AGCGACTCAG GTGCCTGATG GTCGGAGACA AAAGGAATAT<br><br>351 TCAGCGATTT GCCCGAGCTT GCGAGGGTGC TACTTAAGCC TTTAGGGTTT TAAGGTCTGT TTTGTAGAGG<br><br>421 AGCAAACAGC GTTTGCGACA TCCTTTTGTA ATACTGCGGA ACTGACTAAA GTAGTGAGTT ATACACAGGG<br><br>491 CTGGGATCTA TTCTTTTTAT CTTTTTTTAT TCTTTCTTTA TTCTATAAAT TATAACCACT TGAATATAAA<br><br>561 CAAAAAAAAC ACACAAAGGT CTAGCGGAAT TTACAGAGGG TCTAGCAGAA TTTACAAGTT TTCCAGCAAA<br><br>631 GGTCTAGCAG AATTTACAGA TACCCACAAC TCAAAGGAAA AGGACTAGTA ATTATCATTG ACTAGCCCAT<br><br>701 CTCAATTGGT ATAGTGATTA AAATCACCTA GACCAATTGA GATGTATGTC TGAATTAGTT GTTTTCAAAG<br><br>771 CAAATGAACT AGCGATTAGT CGCTATGACT TAACGGAGCA TGAAACCAAG CTAATTTTAT GCTGTGTGGC<br><br>841 ACTACTCAAC CCCACGATTG AAAACCCTAC AAGGAAAGAA CGGACGGTAT CGTTCACTTA TAACCAATAC<br><br>911 GCTCAGATGA TGAACATCAG TAGGGAAAAT GCTTATGGTG TATTAGCTAA AGCAACCAGA GAGCTGATGA<br><br>981 CGAGAACTGT GGAAATCAGG AATCCTTTGG TTAAAGGCTT TGAGATTTTC CAGTGGACAA ACTATGCCAA<br><br>1051 GTTCTCAAGC GAAAAATTAG AATTAGTTTT TAGTGAAGAG ATATTGCCTT ATCTTTTCCA GTTAAAAAAA<br><br>1121 TTCATAAAAT ATAATCTGGA ACATGTTAAG TCTTTTGAAA ACAAATACTC TATGAGGATT TATGAGTGGT |

| | |
|---|---|
| | 1191 TATTAAAAGA ACTAACACAA AAGAAAACTC ACAAGGCAAA TATAGAGATT AGCCTTGATG AATTTAAGTT<br><br>1261 CATGTTAATG CTTGAAAATA ACTACCATGA GTTTAAAAGG CTTAACCAAT GGGTTTTGAA ACCAATAAGT<br><br>1331 AAAGATTTAA ACACTTACAG CAATATGAAA TTGGTGGTTG ATAAGCGAGG CCGCCCGACT GATACGTTGA<br><br>1401 TTTTCCAAGT TGAACTAGAT AGACAAATGG ATCTCGTAAC CGAACTTGAG AACAACCAGA TAAAAATGAA<br><br>1471 TGGTGACAAA ATACCAACAA CCATTACATC AGATTCCTAC CTACATAACG GACTAAGAAA AACACTACAC<br><br>1541 GATGCTTTAA CTGCAAAAAT TCAGCTCACC AGTTTTGAGG CAAAATTTTT GAGTGACATG CAAAGTAAGT<br><br>1611 ATGATCTCAA TGGTTCGTTC TCATGGCTCA CGCAAAAACA ACGAACCACA CTAGAGAACA TACTGGCTAA<br><br>1681 ATACGGAAGG ATCTGAGGTT CTTATGGCTC TTGTATCTAT CAGTGAAGCA TCAAGACTAA CAAACAAAAG<br><br>1751 TAGAACAACT GTTCACCGTT ACATATCAAA GGGAAAACTG TCCATATGCA CAGATGAAAA CGGTGTAAAA<br><br>1821 AAGATAGATA CATCAGAGCT TTTACGAGTT TTTGGTGCAT TCAAAGCTGT TCACCATGAA CAGATCGACA<br><br>1891 ATGTAACAGA TGAACAGCAT GTAACACCTA ATAGAACAGG TGAAACCAGT AAAACAAAGC AACTAGAACA<br><br>1961 TGAAATTGAA CACCTGAGAC AACTTGTTAC AGCTCAACAG TCACACATAG ACAGCCTGAA ACAGGCGATG<br><br>2031 CTGCTTATCG AAGTCTGACG CTCAGTGGAA CGAAAACTCA CGTTAAGGGA TTTTGGTCAT GCATATGAAT<br><br>2101 ATCCTCCTTA GTTCCTATTC CGAAGTTCCT ATTCTCTAGA AAGTATAGGA ACTTCAGAGC GCTTTTGAAG<br><br>2171 CTGGGGTGGG CGAAGAACTC CAGCATGAGA TCCCCGCGCT GGAGGATCAT CCAGCCGGCG TCCCGGAAAA<br><br>2241 CGATTCCGAA GCCCAACCTT TCATAGAAGG CGGCGGTGGA ATCGAAATCT CGTGATGGCA GGTTGGGCGT<br><br>2311 CGCTTGGTCG GTCATTTCGA ACCCCAGAGT CCCGCTTACG CCCCGCCCTG CCACTCATCG CAGTACTGTT<br><br>2381 GTAATTCATT AAGCATTCTG CCGACATGGA AGCCATCACA GACGGCATGA TGAACCTGAA TCGCCAGCGG<br><br>2451 CATCAGCACC TTGTCGCCTT GCGTATAATA TTTGCCCATA GTGAAAACGG GGGCGAAGAA GTTGTCCATA |

| | |
|---|---|
| | 2521 TTGGCCACGT TTAAATCAAA ACTGGTGAAA CTCACCCAGG GATTGGCTGA TACGAAAAAC ATATTCTCAA<br><br>2591 TAAACCCTTT AGGGAAATAG GCCAGGTTTT CACCGTAACA CGCCACATCT TGCGAATATA TGTGTAGAAA<br><br>2661 CTGCCGGAAA TCGTCGTGGT ATTCACTCCA GAGCGATGAA AACGTTTCAG TTTGCTCATG GAAAACGGTG<br><br>2731 TAACAAGGGT GAACACTATC CCATATCACC AGCTCACCGT CTTTCATTGC CATACGGAAC TCCGGATGAG<br><br>2801 CATTCATCAG GCGGGCAAGA ATGTGAATAA AGGCCGGATA AAACTTGTGC TTATTTTTCT TTACGGTCTT<br><br>2871 TAAAAAGGCC GTAATATCCA GCTGAACGGT CTGGTTATAG GTACATTGAG CAACTGACTG AAATGCCTCA<br><br>2941 AAATGTTCTT TACGATGCCA TTGGGATATA TCAACGGTGG TATATCCAGT GATTTTTTTC TCCATGCGAA<br><br>3011 ACGATCCTCA TCCTGTCTCT TGATCAGATC TTGATCCCCT GCGCCATCAG ATCCTTGGCG GCAAGAAAGC<br><br>3081 CATCCAGTTT ACTTTGCAGG GCTTCCCAAC CTTACCAGAG GGCGCCCCAG CTGGCAATTC CGGTTCGCTT<br><br>3151 GCTGTCCATA AAACCGCCCA GTCTAGCTAT CGCCATGTAA GCCCACTGCA AGCTACCTGC TTTCTCTTTG<br><br>3221 CGCTTGCGTT TTCCCTTGTC CAGATAGCCC AGTAGCTGAC ATTCATCCGG GGTCAGCACC GTTTCTGCGG<br><br>3291 ACTGGCTTTC TACGTGTTCC GCTTCCTTTA GCAGCCCTTG CGCCCTGAGT GCTTGCGGCA GCGTGGGGGA<br><br>3361 TCTTGAAGTT CCTATTCCGA AGTTCCTATT CTCTAGAAAG TATAGGAACT TCGAAGCAGC TCCAGCCTAC<br><br>3431 ACGGGAGAGT GTTCACCGAC AAACAACAGA TAAAACAAAA GGCCCAGTCT TCCGACTGAG CCTTTTGTTT<br><br>3501 TATTTGATGT CTGGCAGTTC CCCACTCGCC AGATTTACGA AGTGACGAAT TACTTATCTG GCAGGAGTAT<br><br>3571 CGTCCGTAGT TTCAACTGTT CTGTGCATAC GGCCCTGAAA GACTATTGAT TACGAATATA AGGAAAAACCA<br><br>3641 CTGCTAAGAC GAATTACTTA TCTGGCAGGA GTATCGTCCG TAGTTTCAAC TGTTCTGTGC ATACGGCCCT<br><br>3711 GAAAGACTAT TGATTACGAA TATAAGGAAA ACCACTAGAG ACCTCCTTGG ACCTGGTACG TCGCTAGCGG<br><br>3781 GTAGTGTGAC GATATGTACA CATTATATCA CATCTATTCC AAAATGTCAA CTGAAGGCGC TCCTCATACT |

| | |
|---|---|
| | 3851 GGCTGAATAC ATGACCCGAC ACGTTATCAG AATTTATCAT GTCATGATTG CTTACAATAC TCACATAATT<br><br>3921 GTGAAAATTG TAGAAGCCGC AACATTCAAT TATCAATCAA AATCATGTCC GCGCCAGTTT CGTTCCTCGC<br><br>3991 CGTTGCACTA ATACAGGGTA ATTCCCTCAC CACATAAGGA ATTAATGACG TCAGGTGGCA CTTTTCCACA<br><br>4061 CAGGAGAGCG TTCACCGACA AACAACAGAT AAAACGAAAG GCCCAGTCTT TCGACTGAGC CCTTCGTTTT<br><br>4131 ATTTGATGCC TGGCAGTTCC CAAATGAACG ATTTCTTAGT CGGCGTTATA GTAAGTCACT CTTTTTCAGC<br><br>4201 GGTATTTTAA AGATGAGAAA GCGATGGTCA AGCGTGGTCT GCCTGAAGTC TCAATCAGTC GGAACGGCAA<br><br>4271 TGCTAGCCCA TGCTTANNNN NNNNNNATGG TTAGCGAGCT GATCAAAGAA AACATGCACA TGAAACTGTA<br><br>4341 TATGGAAGGC ACCGTGAATA ACCACCACTT TAAATGTACC AGCGAAGGTG AAGGTAAACC GTATGAAGGC<br><br>4411 ACCCAGACCA TGCGTATTAA AGCAGTTGAA GGTGGTCCGC TGCCGTTTGC ATTTGATATT CTGGCAACCA<br><br>4481 GCTTTATGTA TGGCAGCAAA ACCTTTATTA ACCATACCCA GGGTATCCCG GATTTTTTCA AACAGAGCTT<br><br>4551 TCCGGAAGGT TTTACCTGGG AACGTGTTAC CACCTATGAA GATGGTGGTG TTCTGACCGC AACCCAGGAT<br><br>4621 ACCAGTCTGC AGGATGGTTG TCTGATTTAT AATGTGAAAA TTCGCGGTGT GAACTTTCCG AGCAATGGTC<br><br>4691 CGGTTATGCA GAAAAAAACC CTGGGTTGGG AAGCAAGCAC CGAAACCCTG TATCCGGCAG ATGGTGGTCT<br><br>4761 GGAAGGTCGT GCAGATATGG CACTGAAACT GGTTGGTGGT GGTCATCTGA TTTGCAATCT GAAAACCACC<br><br>4831 TATCGTAGCA AAAAACCGGC AAAAAATCTG AAAATGCCTG GCGTGTATTA TGTTGATCGT CGTCTGGAAC<br><br>4901 GTATTAAAGA GGCAGATAAA GAAACCTATG TGGAACAGCA TGAAGTTGCA GTTGCACGTT ATTGTGATCT<br><br>4971 GCCGAGCAAA CTGGGTCACC GCTGATAACC ATGGGCTAGC GGTTTGAAGG GTATTGGTCG GTCAGTTTCA<br><br>5041 CCTGATTTAC GTAAAAACCC GCTTCGGCGG GTTTTTGCTT TTGGAGGGGC AGAAAGATGA ATGACTGTCC<br><br>5111 TTTTATTGGA GAGGTGGACA AGTGGCATCA GAGTTCACTC TTAATTCTGA ACATACCCGT CTTTTTCGCC |

| | |
|---|---|
| | 5181 TCTTTTACGT GATTAACTCC AGCGCTGGCG GCGGTTTTTA AAGGACAAAG ACTCCGGTAT TCAGACATGA<br><br>5251 CAACAAATTA CCAGGGTTTG GCTGCCGGAC ATAAAATTTT GGTTTACAGC AATTTATATA TTCCAGTCGG<br><br>5321 GAAACCTGTC GTGCCAGCTG CATTAATGAA TCGGCCAACG CGAATTCCCG ACA |

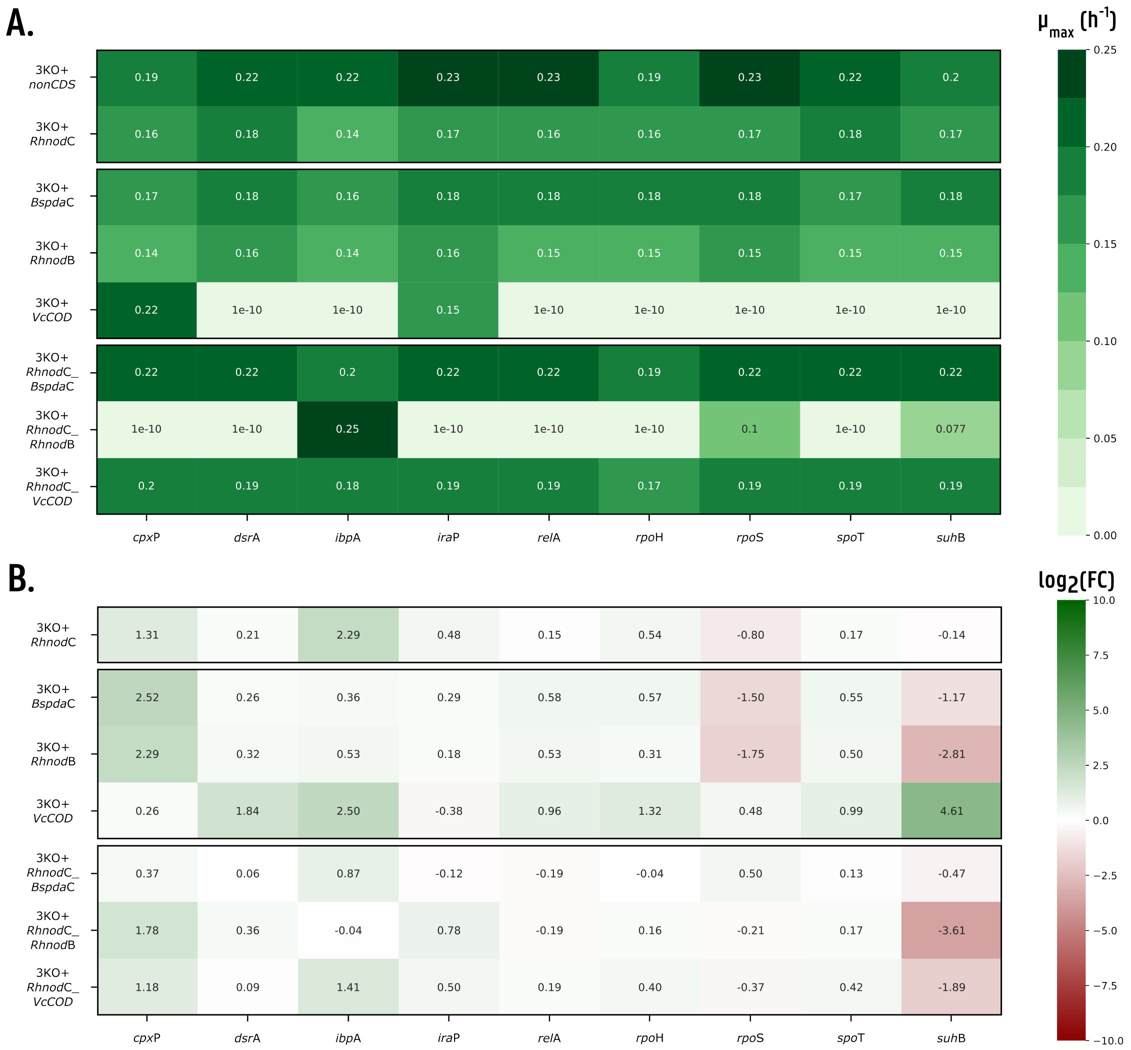

**Supplementary Figure 1:** Heatmaps of the results for the stress promoter assay for the expression of RhNodC and/or different chitin deacetylases. **A.** Growth rate (μmax ($h^{-1}$)) of the strains. **B.** $Log_2$(FC) of the corrected fluorescence of the stress promoters in the stationary phase compared to the control strain 3KO+*nonCDS*+pInd and **C.** in the exponential phase. Biologically significant results ($Log_2$(FC) > 1) have a higher color intensity. Details of the strains, plasmids and stress promoters used can be found in Table 4, Supplementary Table 8 and Table 2, respectively. $Log_2$(FC) = $log_2$ fold change.

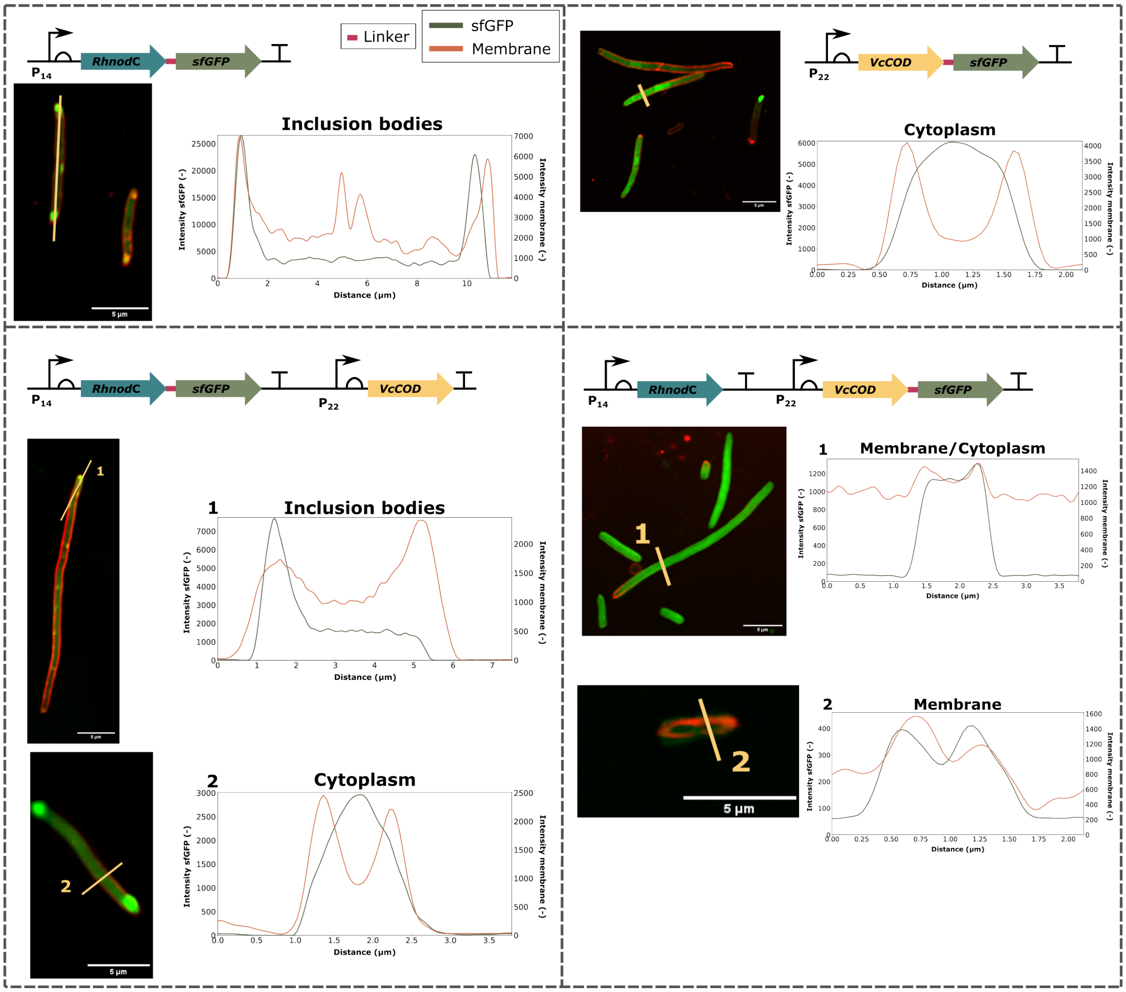

**Supplementary Figure 2:** Confocal light scanning microscopy images for *Rh*NodC and *Vc*COD with intensity cross-section profiles for both the red FM4-64 membrane dye (orange) and sfGFP (green). The cross-sections analysed are indicated in yellow in the images. *Rh*NodC = chitin oligosaccharide synthase from *Rhizobium* sp. GRH2; *Vc*COD = chitin deacetylase from *Vibrio cholerae*; sfGFP = superfolder green fluorescent protein. Gene structures are displayed according to the SBOL guidelines [1].

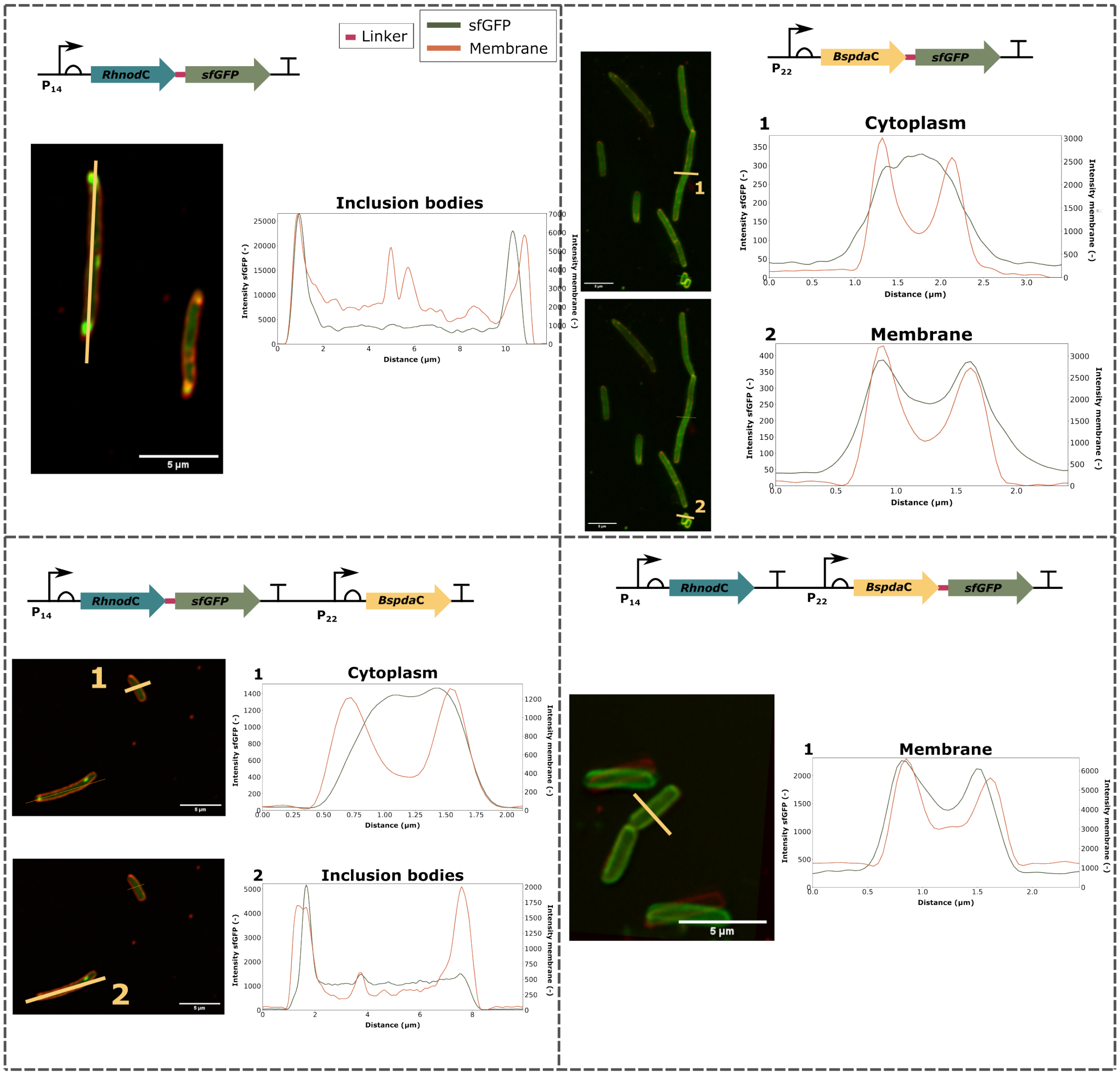

**Supplementary Figure 3:** Confocal light scanning microscopy images for *Rh*NodC and *Bs*PdaC with intensity cross-section profiles for both the red FM4-64 membrane dye (orange) and sfGFP (green). The cross-sections analyzed are indicated in yellow in the images. *Rh*NodC = chitin oligosaccharide synthase from *Rhizobium* sp. GRH2; *Bs*PdaC= chitin deacetylase from *Bacillus subtilis*; sfGFP = superfolder green fluorescent protein. Gene structures are displayed according to the SBOL guidelines [1].

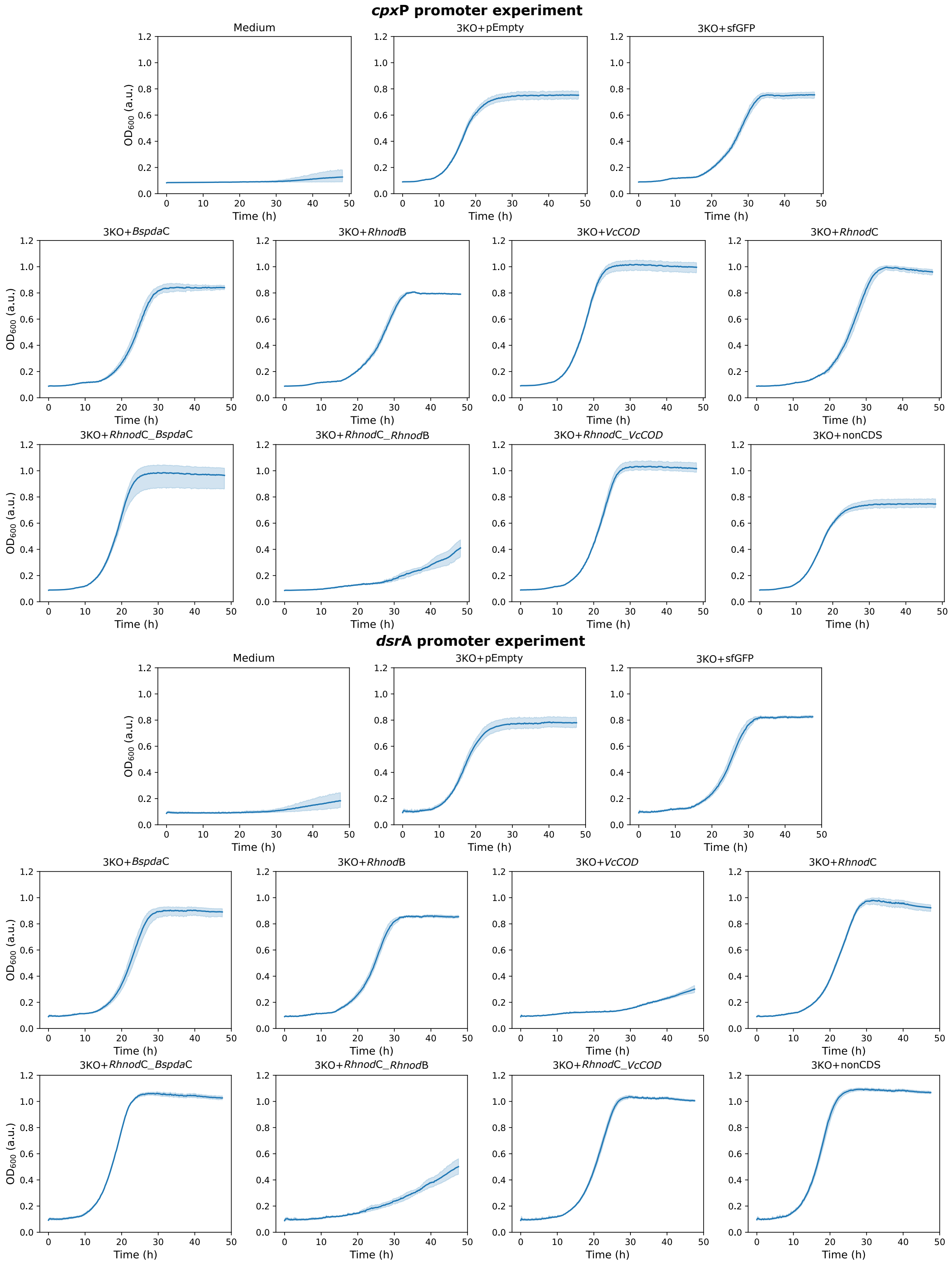

**Supplementary Figure 4: Growth curves obtained during the stress promoter assay.** *cpx*P- and *dsr*A-promoter experiment curves are shown for the medium, control strains and production strains.

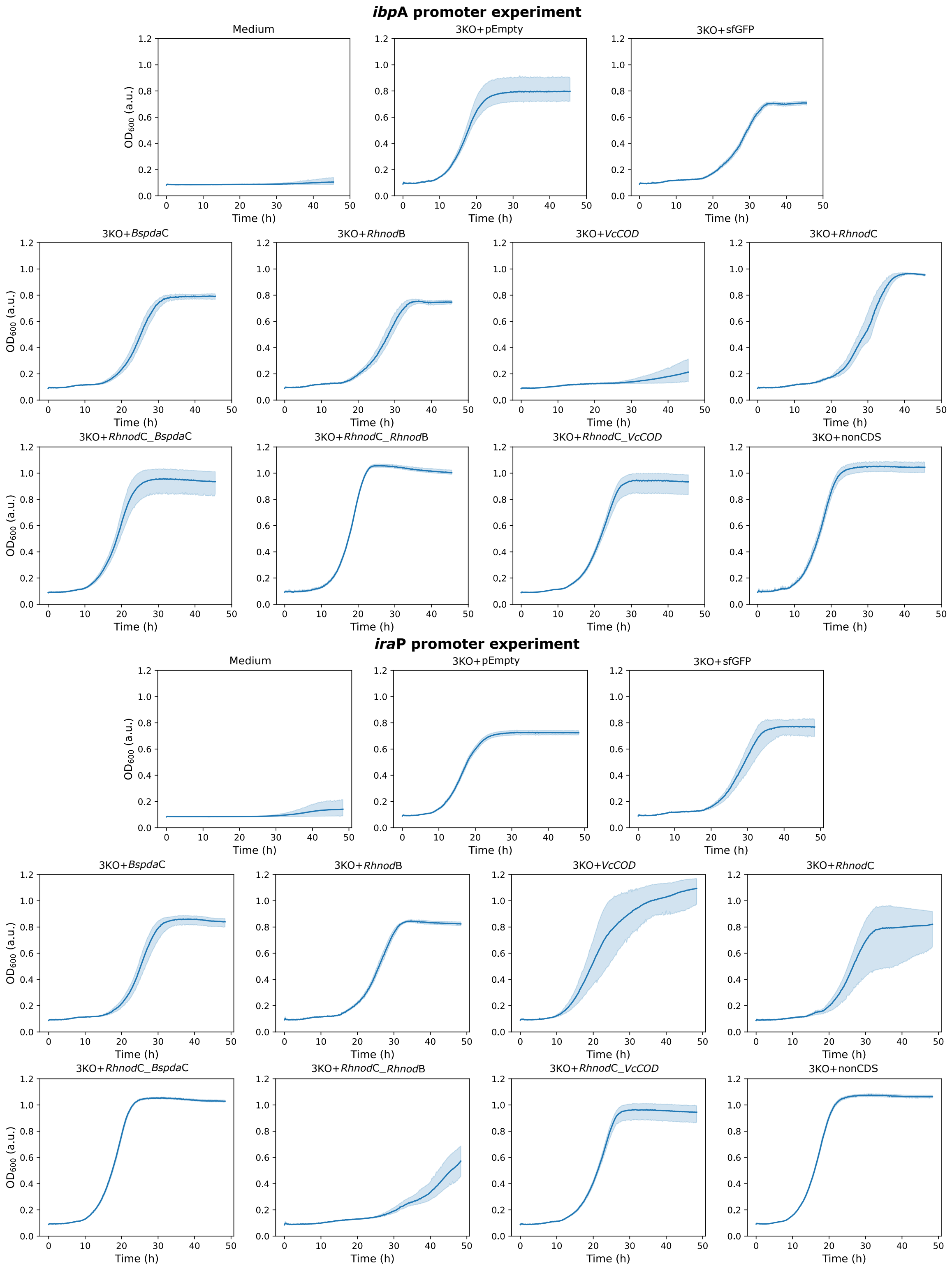

**Supplementary Figure 4: Growth curves obtained during the stress promoter assay.** *ibp*A- and *ira*P-promoter experiment curves are shown for the medium, control strains and production strains (continued).

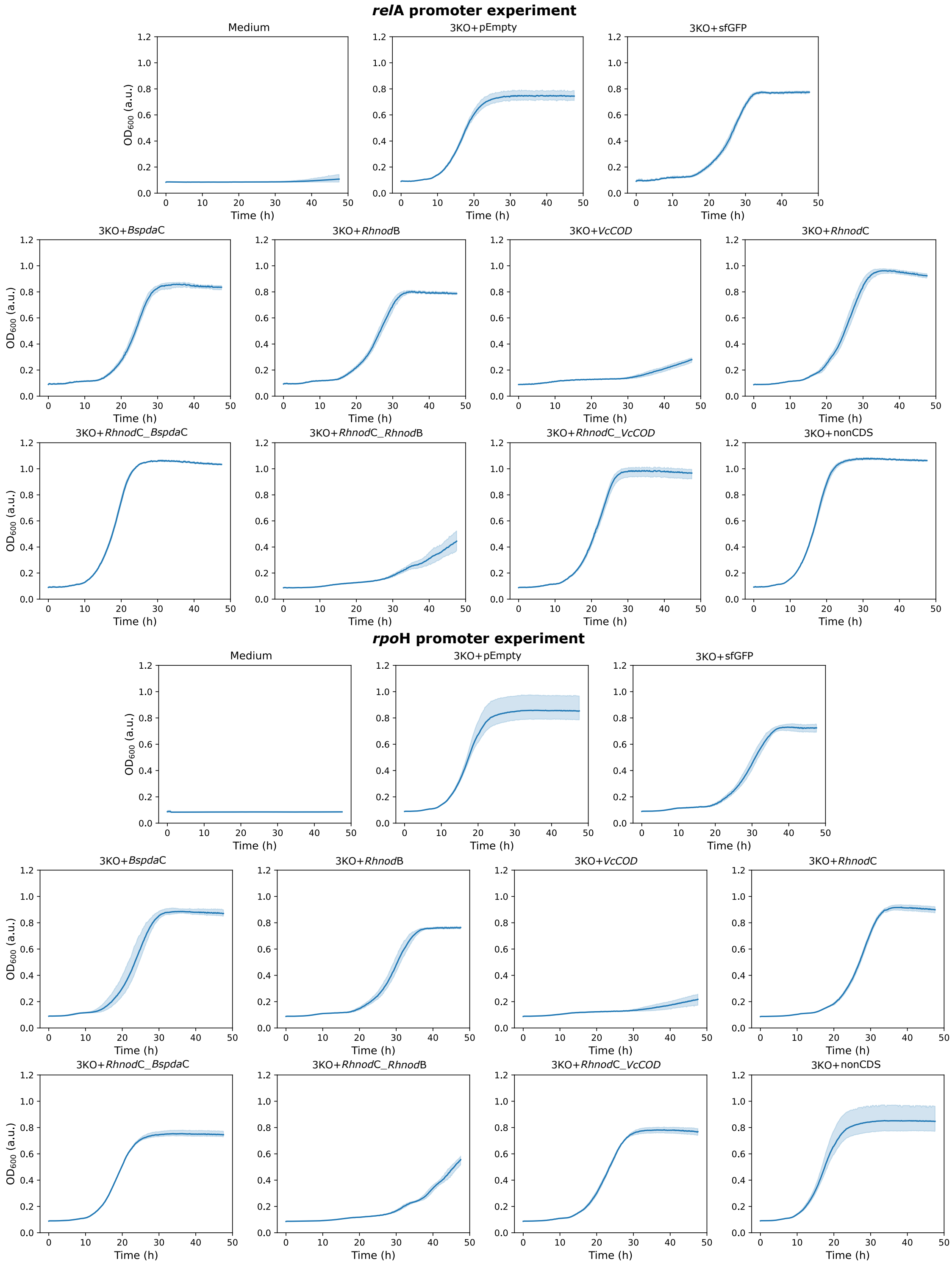

**Supplementary Figure 4: Growth curves obtained during the stress promoter assay.** *rel*A- and *rpo*H-promoter experiment curves are shown for the medium, control strains and production strains (continued).

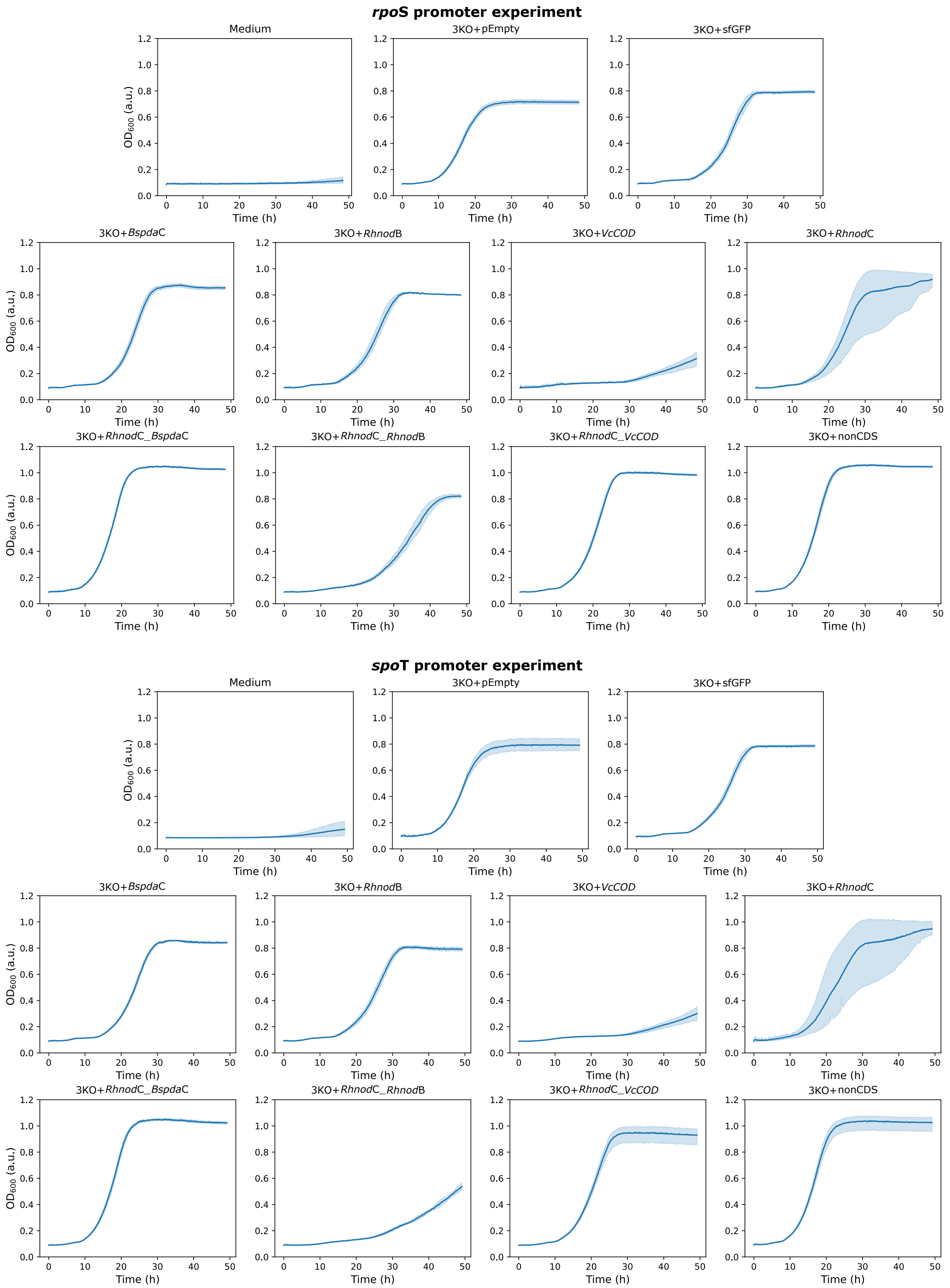

**Supplementary Figure 4: Growth curves obtained during the stress promoter assay.** *rpo*S- and *spo*T-promoter experiment curves are shown for the medium, control strains and production strains (continued).

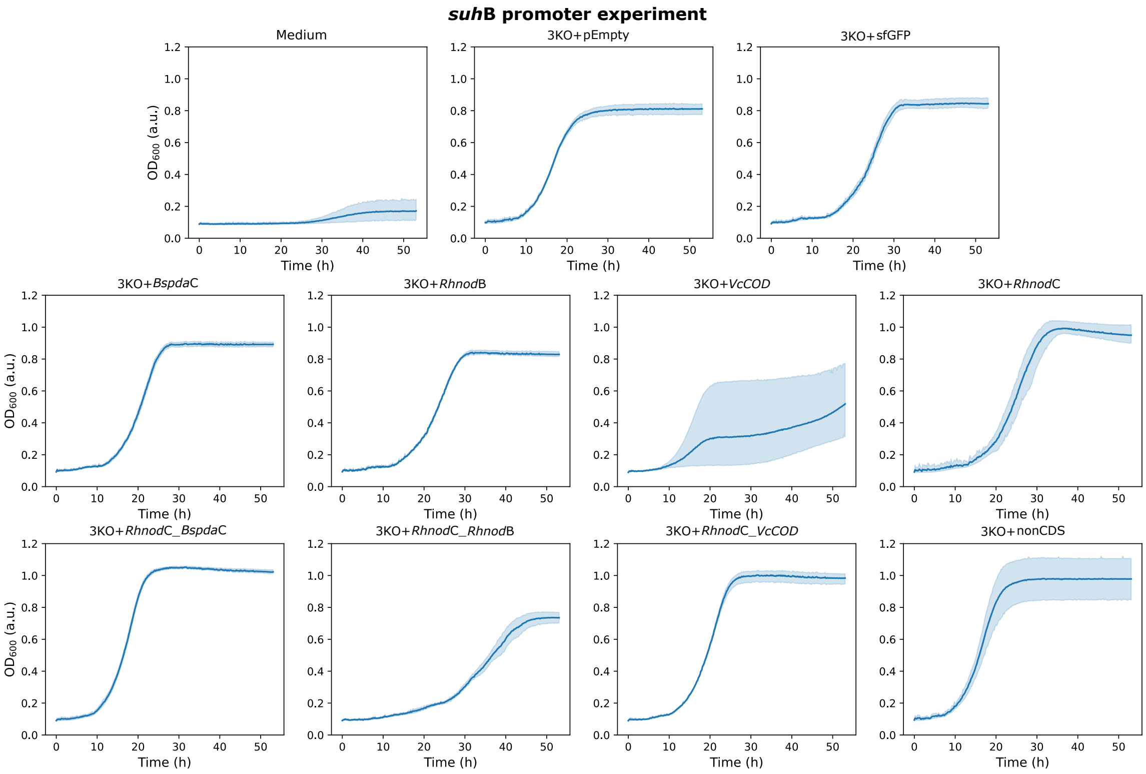

**Supplementary Figure 4: Growth curves obtained during the stress promoter assay.** The *suh*B promoter experiment curves are shown for the medium, control strains and production strains (continued).
